# Basal ganglia output dynamically controls skilled forelimb kinematics in real time

**DOI:** 10.64898/2026.03.09.710687

**Authors:** Shaolin Ruan, Henry H. Yin

**Affiliations:** Department of Neurobiology, Duke University School of Medicine, Durham, NC, 6 27708, USA; Department of Psychology and Neuroscience, Duke University, Durham, NC, 7 27708, USA

**Keywords:** basal ganglia, substantia nigra pars reticulata, motor control, kinematics, reaching

## Abstract

The substantia nigra pars reticulata (SNr), the principal output nucleus of the basal ganglia, is traditionally viewed as a binary gate that permits actions through disinhibition. However, this framework fails to explain the fluid, high-dimensional control required for skilled behavior. Using high-resolution 3D kinematic tracking and automated behavioral classification, we show that SNr population activity paradoxically increases during skilled forelimb reaching and scales with motif-specific kinematics. Systematic optogenetic and chemogenetic perturbations reveal that the SNr bidirectionally controls movement vigor and online kinematics. Remarkably, a transient 12.5 ms pause is sufficient to abort a motor program, while a 12.5 ms burst enhances velocity and reshapes trajectories without altering sequence identity. Calcium imaging at single-neuron resolution further confirms that endogenous SNr dynamics represent real-time 3D kinematics. These findings demonstrate that SNr output does not merely gate movement initiation but continuously regulates the stability and kinematic evolution of skilled actions, expanding the functional framework of basal ganglia output from binary selection to the real-time dynamical control of motor execution.

**Highlights:**

- SNr activity paradoxically increases and scales with motor motif kinematics
- Bidirectional SNr control of action vigor for skilled forelimb reaching
- Brief SNr pauses or bursts reshape online motor execution
- The SNr acts as a continuous controller for real-time motor execution

## Introduction

The execution of skilled forelimb movements exemplifies the sophistication of mammalian motor control, requiring the brain to select the appropriate action while precisely calibrating its vigor and controlling its kinematics in real-time. While this orchestration involves a distributed network of brain regions^1–8^, the basal ganglia (BG) are essential^9,10,3,11–16^, with the substantia nigra pars reticulata (SNr) serving as the primary output nucleus^17,18^. The SNr sends inhibitory projections to the thalamus and brainstem^19–24^, but the functional role of this output remains unclear.

Classic models of BG function emphasize a gating mechanism. Inhibitory striatal projections to the SNr can produce a pause in the tonically active, inhibitory SNr neurons to permit movement via disinhibition^25,26^. While this model provides a coherent framework for action selection^27–31^, it struggles to reconcile the binary nature of gating with the continuous, high-dimensional control required for the execution of skilled motor sequences. Recent evidence suggests that the striatum, the primary input nuclei of the BG^32^, exerts significant influence over movement vigor^33–35^ and online kinematics^36,37,11^, yet it remains unclear how the BG shapes movement at the output stage. Does the SNr merely relay a binary go/no-go signal, as assumed in traditional models, or does it contribute to the vigor and the real-time kinematic control of movement.

This gap is further accentuated by a paradox in the literature on the SNr: contrary to the canonical disinhibition model^25,26^, some recent observations reveal that many SNr neurons increase their firing during behavior^16,19,38–42^, often outnumbering those that decrease^16,41,42^. This paradoxical recruitment suggests that the traditional model is incomplete at best. Importantly, it remains to be determined whether this active recruitment is a mere byproduct of circuit dynamics or a functional requirement for the execution of skilled behavior.

To address these questions, we developed a high-resolution 3D tracking paradigm and a GRU-based classifier to decompose the reaching sequence into a series of stereotyped motor motifs. This high-vigor, and online kinematics at millisecond temporal resolution. In this study, we provide evidence that fundamentally challenges the canonical disinhibition model. We reveal a recruitment of the SNr population that serves as a continuous, real-time drive for execution of skilled forelimb movements. By momentarily pausing and augmenting SNr activity during specific behavioral windows corresponding to individual action primitives, we demonstrate that SNr can continuously shapes the spatial and temporal parameters of movement. Together, these findings redefine the SNr not as a simple binary gate, but as a dynamic orchestrator of the fine-grained kinematics in skilled behavior.

## Results

### Paradoxical increase in SNr population activity underlying stereotyped forelimb reaching action sequence

To investigate the contribution of basal ganglia output to the execution of skilled forelimb movements, we developed a high-resolution water-reaching paradigm for freely moving mice. This setup leveraged two orthogonal, synchronized high-speed cameras to permit the 3D kinematic reconstruction of forelimb trajectories (Figure 1A). To achieve unbiased, high-throughput behavioral analysis, we implemented a Gated Recurrent Unit (GRU)-based classifier^43^ that utilized augmented body-part coordinates to automatically segment reaching sequences into discrete motor motifs. These segmented labels and their associated kinematics served as the foundation for all subsequent motif-specific analyses (Figure S1A).

**Figure 1.**
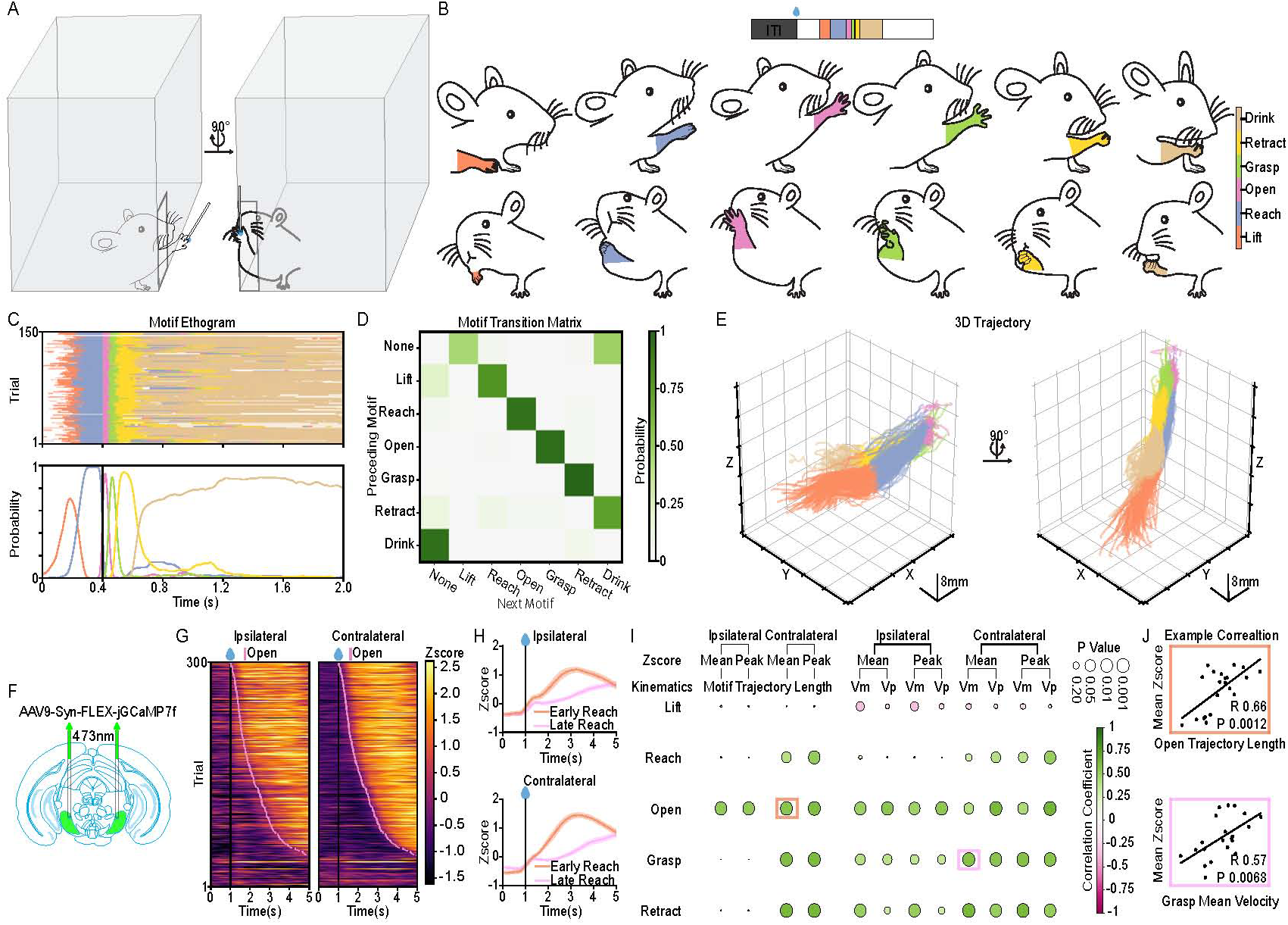
Increase of SNr population activity underlying stereotyped forelimb reaching action sequence. **(A)** Schematic of the free-moving behavioral setup utilizing two perpendicular high-speed cameras for simultaneous recording and 3D reconstruction of forelimb reaching kinematics. **(B)** Decomposition of the reaching sequence into six distinct, stereotyped action motifs (**Lift**, **Reach**, **Open**, **Grasp**, **Retract**, **Drink**). Top and bottom rows show orthogonal views of the posture for each motif. Colors correspond to specific motifs throughout the figure (color legend inset). Trial-based task structure illustrated above. **(C)** Representative motif ethogram (top) and corresponding motif probability (bottom) from an example session (150 trials), aligned to the first **Open** motif. **(D)** Motif transition matrix from the representative session in (C). Each color block represents the transition probability from the motif on the y-axis to the motif on the x-axis. **(E)** Reconstructed 3D trajectories of the forelimb for all trials in the representative session in (C), color-coded by motif. **(F)** Schematic of the fiber photometry strategy. AAV9-Syn-FLEX-jGCaMP7f was injected into the SNr to record populational calcium dynamics in SNr GABAergic neurons during behavior. **(G)** Representative heatmaps of Z-scored SNr calcium signals from ipsilateral (left) and contralateral (right) hemispheres, aligned to water delivery and sorted by the onset of the first **Open** motif (pink line). Error bars represent mean ± SEM. **(H)** Average Z-scored SNr activity aligned to water delivery, stratified as early reaching trials and late reaching trials by reaction time (top 30% vs. bottom 30%). n = 21 sessions (2100 trials), 7 mice. **(I)** Correlation matrix summarizing the relationship between motif-specific kinematics (Trajectory Travelled Length, Mean/Peak Velocity) and motif specific SNr Z-scores. Circle size denotes statistical significance (p-value). Color intensity indicates the correlation coefficient. Spearman’s rank correlation coefficient was used (Supplementary Table 1). n = 21 sessions (2100 trials), 7 mice. Refer to Table 1 for summary statistics. **(J)** Representative scatter plots showing significant positive correlations between SNr activity and specific kinematic parameters: **Open** motif trajectory length (top) and **Grasp** motif mean velocity (bottom). Colored outlines in (I) indicate the specific correlations displayed here. n = 21 sessions (2100 trials), 7 mice. Pearson correlation coefficient was used.

The water-reaching sequence can be consistently decomposed into six distinct, stereotyped action motifs: Lift, Reach, Open, Grasp, Retract, and Drink (Figure 1B and Video S1). Each motif was defined by a unique and reproducible kinematic signature (Figure S1B–M). For instance, the Open and Grasp phases were characterized by minimum proximity to the water spout (Figure S1F) and a maximum increase in inter-digit distance (Figure S1G). Furthermore, inter-digit velocity peaked specifically during these dexterous phases, contrasting with the more ballistic profiles observed during the Reach and Retract segments (Figure S1K, M). The progression through this motor program was highly organized and reliable, as evidenced by the structured behavioral ethograms (Figure 1C) and the high probability along the superdiagonal in the motif transition matrix (Figure 1D). Consistent with this modular organization, 3D trajectory reconstructions revealed a remarkably stereotyped execution of the reaching sequence across trials (Figure 1E and Video S2).

To investigate the recruitment of the SNr during skilled forelimb tasks at the population level, we expressed the genetically encoded calcium indicator jGCaMP7f in the SNr and performed fiber photometry during the water-reaching task (Figure 1F). Contrary to the classical disinhibition model, which predicts a pause in SNr output during movement, we observed a robust increase in both ipsilateral and contralateral SNr population activity during the reaching sequence (Figure 1G). To determine if this activity was coupled to the initiation of the motor program, we stratified trials by reaction time (fastest 30% vs. slowest 30%). In both hemispheres, the peak of SNr calcium dynamics tracked the temporal shift in behavior, peaking significantly earlier in fast-reaction trials compared to slow-reaction trials (Figure 1H).

We next examined whether these population dynamics encoded specific kinematic parameters of the forelimb. We performed a correlation analysis between SNr activity and three key kinematic features: motif trajectory length, mean velocity, and peak velocity. SNr activity was positively correlated with these motif specific kinematics parameters across several motifs, from the Reach through the Retract motif. Notably, these correlations seem to be more pronounced in the contralateral hemisphere (Figure 1I), suggesting that the encoding of skilled movements may be lateralized. Representative correlation plots (Figure 1J) further illustrate that SNr activity does not merely signal movement onset, but rather scales with the physical execution of the motor motif.

Collectively, these data demonstrate that, at least at the population level as measured by photometry recording, an increase, rather than a pause, in SNr neurons is associated with the execution of stereotyped reaching, scaling with the kinematics of the action sequence.

### SNr activation is required for action selection and vigor control of dexterous reaching behavior

The observed recruitment of the SNr population during reaching suggests that these dynamics are not merely correlative, but are functionally required for motor execution. To test this hypothesis, we employed optogenetic inhibition using the inhibitory opsin GtACR2, delivering photo-stimulation (473 nm, 4 s) at water delivery to silence SNr activity throughout the reaching sequence (Figure 2A).

**Figure 2.**
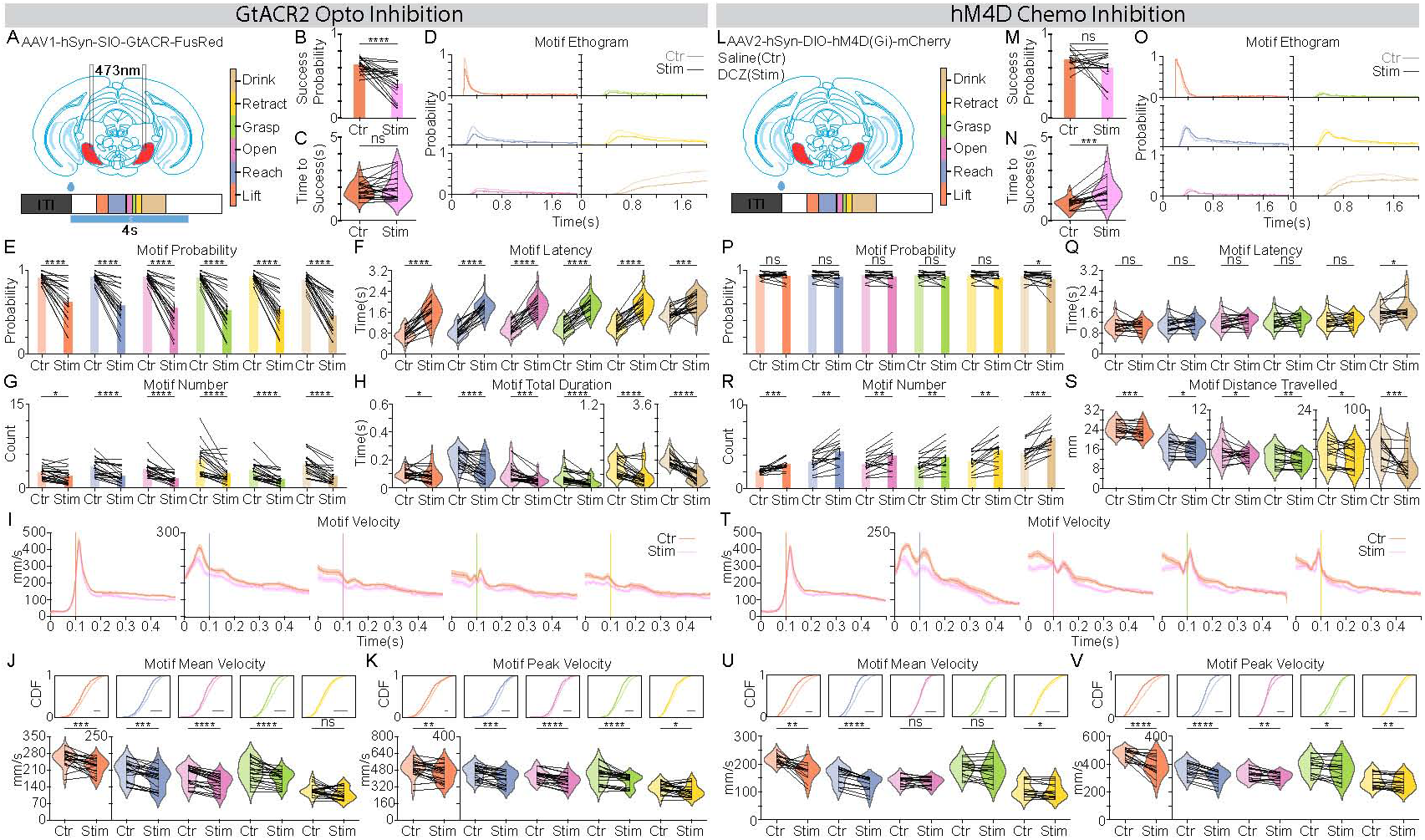
SNr inhibition suppresses forelimb skilled reaching behavior. **(A)** Schematics of the experimental strategies. Optogenetic inhibition: AAV1-hSyn-DIO-GtACR2-FusRed was injected into the SNr, and photoinhibition (473nm, 4s) was delivered starting at water delivery. The motif color map (inset) applies to all panels. **(B and C)** Comparison of reach success parameters: **(B)** Probability of success and **(C)** Latency to success relative to reach onset. **(D)** Superimposed motif ethograms aligned to the first **Lift** motif, comparing inhibition (saturated colors) and control (lighter colors) trials. **(E and F)** Comparison of reach initiation dynamics: **(E)** Motif occurrence probability and **(F)** motif latency. **(G)** Quantification of motif count per trial. **(H)** Quantification of total motif duration. **(I)** Forelimb velocity profiles (mean ± SEM) aligned to the onset of **Lift**, **Reach**, **Open**, **Grasp**, and **Retract** motifs (left to right). Orange lines: control; Purple lines: stimulation. **(J and K)** Comparison of motif Mean Velocity **(J)** and Peak Velocity **(K)**. Top panels show the Cumulative Distribution Function (CDF) plots of all trials; bottom panels show the violin plots of all sessions. **(L)** Schematics of the experimental strategies. Chemogenetic inhibition: AAV2-hSyn-DIO-hM4D(Gi)-mCherry was injected into the SNr; Deschloroclozapine (DCZ) served as the agonist, with saline as the control. **(M and N)** as in **(B and C)** respectively. **(O)** as in **(D)**. **(P and Q)** as in **(E and F)** respectively. **(R)** as in **(G)**. **(S)** Quantification of motif distance travelled. **(T)** as in **(I)**. **(U and V)** as in **(J and K)** respectively. Left panel (GtACR2 Optogenetic inhibition), n = 2,687 control trials and 1,094 stimulation trials from 21 sessions, 7 mice. Right panel (hM4D Chemogenetic inhibition), n = 1,125 control trials and 1,125 stimulation trials from n= 15 sessions, 5 mice. Data in bar plot represent mean ± SEM. Dots in the bar and violin plots represent individual session means. Violin plots display the full data distribution density; internal box plots represent the median (center line) and the interquartile range. Wilcoxon Signed-Rank Test was used. ns no significance, *p < 0.05, **p < 0.01, ***p < 0.001, ****p < 0.0001.

In striking contrast to classical disinhibition models, which would predict that silencing the SNr might facilitate movement, optogenetic inhibition significantly suppressed reaching, as shown in the reduced success probability (Figure 2B). While the time to success remained unchanged in the sparse subset of successful trials (Figure 2C), the overall behavioral output was severely stunted (Figure 2D-H and Video S3, 4). Motif ethograms (Figure 2D) and quantification of motif initiation-related parameters (Figure 2E–H). revealed a global suppression across all motor motifs. Specifically, SNr inhibition led to a significant decrease in motif occurrence probability (Figure 2E), an increase in latency (Figure 2F), and a reduction in both motif count (Figure 2G) and total duration per trial (Figure 2H).

Beyond action selection, we examined whether the SNr also modulates motor vigor by analyzing forelimb velocity. We found that silencing the SNr resulted in a marked reduction in forelimb velocity across the Lift, Reach, Open, Grasp, and Retract motifs (Figure 2I and Video S4). Quantifying these effects revealed significant decreases in both mean and peak velocities for nearly all motor motifs (Figure 2J–K), suggesting a role of SNr activation in maintaining forelimb vigor.

To further validate the specificity of these effects and demonstrate that reaching behavior is not just being interrupted by the laser inhibition but is functionally titrated by the level of SNr activation, we performed a dose-response analysis by varying the intensity of SNr photoinhibition (1.25 mW, 2.5 mW, 5 mW, and 10 mW; Figure S2A). We observed a robust, intensity-dependent suppression of reaching: higher laser power resulted in lower reach probabilities during the stimulation period (Figure S2B) and a correspondingly higher rebound reaching probability immediately following laser offset (Figure S2C). This titration was further reflected in the latency to reach, which scaled linearly with inhibition intensity (Figure S2D). Moreover, analysis of motif transition dynamics revealed that increasing inhibition intensity progressively suppressed reaching sequence (Figure S2E), characterized by a reduction in the probability of forward-sequence transitions (along the superdiagonal) and a reciprocal increase in the probability of no-reach states (leftmost column; Figure S2F). Finally, by focusing on lower intensities (1.25 mW and 2.5 mW) where movement was not entirely abolished, we confirmed that motor vigor is also titratable. Greater SNr inhibition led to a gradual, intensity-dependent reduction in both mean and peak forelimb velocities (Figure S2G–I).

One potential concern with focal optogenetic inhibition in an interconnected GABAergic nucleus like the SNr is that silencing a localized population could paradoxically activate distant SNr neurons via the relief of local inhibitory axonal collaterals^44^. To control for this and achieve a more global silencing of the SNr, we employed an alternative approach using chemogenetics. We expressed the inhibitory DREADD hM4D(Gi) in the SNr and administered the potent agonist Deschloroclozapine (DCZ) to inhibit the population more broadly (Figure 2L). While chemogenetic inhibition yielded a less pronounced effect on action selection (Figure 2M–S and Video S3-4), compared to high-intensity photoinhibition, its impact on motor vigor was even more robust (Figure 2T-V). Specifically, we observed no change in overall success probability (Figure 2M), but a significant increase in the time to success (Figure 2N). While motif probability and latency remained largely unaffected (Figure 2O–Q), we observed an increase in the number of motifs per trial accompanied by a significant reduction in the distance traveled for each motif (Figure 2R, S). This indicates that the individual motor motifs were hypometric, necessitating a higher number of reaching attempts to successfully reach water. Consistently, chemogenetic silencing resulted in a marked reduction in forelimb velocity across the Lift, Reach, Open, Grasp, and Retract motifs (Figure 2T–V). These effects on action selection and kinematic vigor closely mirrored those seen with low-intensity photoinhibition (Figure S2), reflecting a more moderate but widespread neural suppression. Ultimately, these chemogenetic results confirm that the suppression of skilled reaching stems from the inhibition of SNr output rather than unintended collateral activation.

Collectively, these data demonstrate that rather than facilitating movement through a pause of inhibition, as suggested by classical models, the activation of the SNr is required for action selection and vigor control of dexterous reaching behavior.

### Post-inhibitory rebound reaches exhibit enhanced vigor

The observed recruitment of the SNr population during reaching, coupled with the suppressive effects of its inhibition, suggests that SNr activation is a key driver of dexterous motor execution. Given that higher photoinhibition intensities triggered rebound reach robustly upon laser offset (Figure S2C), and since prolonged photoinhibition is frequently associated with a compensatory burst of rebound neural activity, we hypothesized that this post-inhibitory rebound neural activity might actively promote subsequent reach. To test this, we silenced the SNr using GtACR2 (473 nm, 5 mW) for randomly interleaved durations of 1 s, 2 s, 3 s, or 4 s (Figure S3A). This protocol reliably suppressed reaching during the laser window and triggered immediate rebound reaching upon offset, regardless of the prior inhibition duration (Figure S3B–D). This temporal precision allowed us to isolate and characterize the specific kinematic signatures of motor behavior driven by a post-inhibitory state. While rebound reaches maintained a normal motif progression (Figure S3E) and achieved success probabilities comparable to control trials (Figure S3F), their latency to success was significantly reduced in rebound trials (Figure S3G), a result that stands in direct opposition to the increased latency observed during SNr inhibition (Figure 2N). Further motif ethogram analysis revealed that these rebound reaches were more vigorous than their control counterparts (Figure S3H). Moreover, we observed a significant increase in forelimb velocity during the initial phases of the sequence, specifically during the Lift, Reach, and Open motifs (Figure S3I–K). Collectively, these results demonstrate that the neural rebound following SNr inhibition drives rebound reaching behavior with enhanced kinematic signatures. This potentiation of motor vigor provides further causal evidence that SNr activation, rather than inhibiting movement, acts as a potent promoter of skilled forelimb reaching.

### SNr activation is a critical, frequency-dependent driver of the kinematic vigor for dexterous reaching

To directly test the hypothesis that SNr activation promotes rather than suppresses skilled motor execution, we employed parallel gain-of-function strategies. We performed focal optogenetic activation using ChR2 (473 nm, 4 s starting at onset of water delivery) and, to account for potential artifacts of local collateral-mediated inhibition, we utilized a more global chemogenetic approach expressing the excitatory DREADD hM3D(Gq) in SNr (Figure 3A, L).

**Figure 3.**
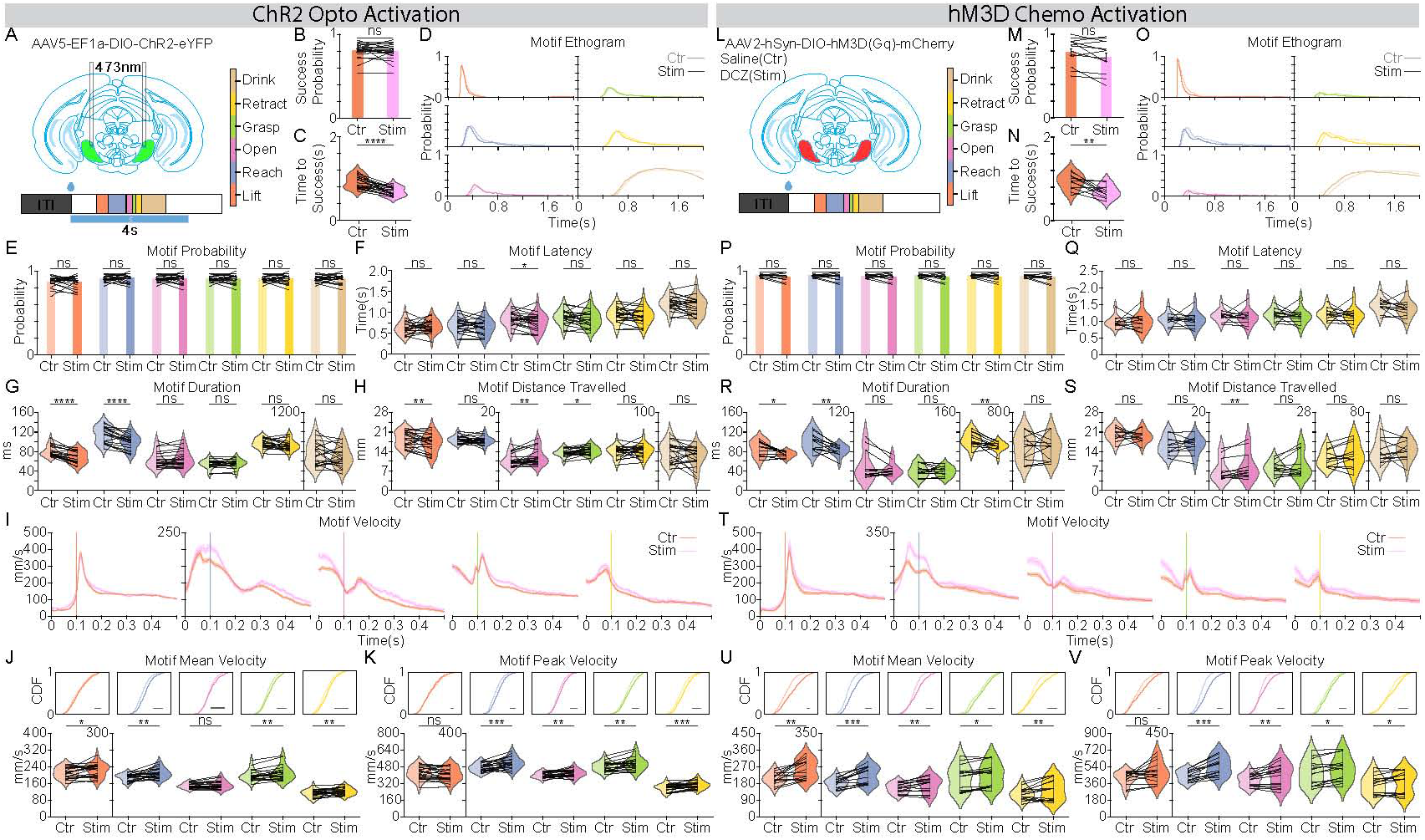
SNr activation promotes forelimb skilled reaching behavior. **(A)** Schematics of the experimental strategies. Optogenetic activation: AAV5-EF1a-DIO-ChR2-eYFP was injected into the SNr, and photoactivation (473nm, 4s) was delivered starting at water delivery. The motif color map (inset) applies to all panels. **(B and C)** Comparison of reach success parameters: **(B)** Probability of success and **(C)** Latency to success relative to reach onset. **(D)** Superimposed motif ethograms aligned to the first **Lift** motif, comparing activation (saturated colors) and control (lighter colors) trials. **(E and F)** Comparison of reach initiation dynamics: **(E)** Motif occurrence probability and **(F)** motif latency. **(G)** Quantification of motif duration. **(H)** Quantification of total motif duration. **(I)** Forelimb velocity profiles (mean ± SEM) aligned to the onset of **Lift**, **Reach**, **Open**, **Grasp**, and **Retract** motifs (left to right). Orange lines: control; Purple lines: stimulation. **(J and K)** Comparison of motif Mean Velocity **(J)** and Peak Velocity **(K)**. Top panels show the Cumulative Distribution Function (CDF) plots of all trials; bottom panels show the violin plots of all sessions. **(L)** Schematics of the experimental strategies. Chemogenetic activation: AAV2-hSyn-DIO-hM3D(Gq)-mCherry was injected into the SNr; Deschloroclozapine (DCZ) served as the agonist, with saline as the control. **(M and N)** as in **(B and C)** respectively. **(O)** as in **(D)**. **(P and Q)** as in **(E and F)** respectively. **(R)** as in **(G)**. **(S)** as in **(H)**. **(T)** as in **(I)**. **(U and V)** as in **(J and K)** respectively. Left panel (ChR2 Optogenetic inhibition), n = 1,824 control trials and 691 stimulation trials from 21 sessions, 4 mice. Right panel (hM3D Chemogenetic inhibition), n = 900 control trials and 900 stimulation trials from n= 12 sessions, 4 mice. Data in bar plot represent mean ± SEM. Dots in the bar and violin plots represent individual session means. Violin plots display the full data distribution density; internal box plots represent the median (center line) and the interquartile range. Wilcoxon Signed-Rank Test was used. ns no significance, *p < 0.05, **p < 0.01, ***p < 0.001, ****p < 0.0001.

Overall, we observed consistent promoting effects across both optogenetic and chemogenetic activation (Video S3-4). Neither focal nor global activation impaired behavioral performance with success probability unaffected by activation (Figure 3B, M), and motif occurrence probabilities and latencies were largely unaffected (Figure 3E, F and P, Q). These findings contradict a recent report suggesting that SNr activation impairs reaching^16^, instead pointing toward an opposite role in skilled forelimb movement. Consistent with this, we observed a significant reduction in the latency to success (Figure 3C, N) and a pronounced leftward shift in the behavioral ethograms (Figure 3D, O), indicating that SNr activation accelerates the progression of the motor program. Detailed kinematic quantification revealed that this enhancement was characterized by motif-specific modulations of duration, distance travelled, and velocity. During the ballistic phases of the sequence (Lift, Reach, and Retract), SNr activation significantly increased movement velocity (Figure I-K and T-V) while simultaneously shortening motif duration (Figure 3G, R). As a result, the total distance traveled during these segments remained relatively stable (Figure 3H, S). In contrast, during the more dexterous middle phases (Open and Grasp), the increase in velocity (Figure 3G, R) was not accompanied by a reduction in duration (Figure I–K and T–V). Consequently, these motifs exhibited a significant increase in trajectory length (Figure 3H, S), driven by the sustained increase in velocity.

To further validate the specificity of these effects and demonstrate that reaching behavior is functionally titrated by the level of SNr activation, we performed a dose-response analysis by varying the frequency of SNr photoactivation (20 Hz and 40 Hz; Figure S4A). Consistent with our main findings, neither reach probability nor the latency to reach were affected by either frequency (Figure S4B–D). Intriguingly, we even observed a slight but significant improvement on success probability during 20 Hz stimulation (Figure S4E). Both activation frequencies produced a comparable reduction in the latency to success (Figure S4F) and a pronounced leftward shift in the behavioral ethograms (Figure S4H), further establishing that SNr activation accelerates motor program progression. Finally, detailed kinematic quantification confirmed that movement vigor can be titrated. Greater SNr activation frequencies led to a gradual, frequency-dependent increase in both mean and peak forelimb velocities across motifs, from the Reach through the Retract segments (Figure S4I–K).

Collectively, these data provide causal evidence that SNr activation, contrary to classical disinhibition theory and recent findings^16^, acts as a motif-specific driver of motor vigor, where activity levels causally scale with the kinematic vigor of skilled forelimb reaching.

### SNr is not involved in the trial-to-trial refinement and reinforcement of forelimb dexterous reaching behavior

While our results establish that SNr activity actively modulates the execution of reaching through action selection and vigor control, it remains unclear how it affects future behavior, e.g. reinforcement of the motor program. Given that reaching is a learned behavior driven by exploration and reinforcement^45^ and that the basal ganglia output is involvement in reinforcing^46^ and punishment^47^, we tested whether SNr optogenetic perturbations leave a functional reinforcement or punishment signal on subsequent motor performance. To investigate this, we performed a trial-history analysis comparing control trials immediately following a photostimulation trial (Post Stim) to those following a control trial (Post Ctr). We reasoned that if the SNr were involved in the refinement and reinforcement of the motor program, the behavioral profile of the Post Stim trial would deviate from that of the Post Ctr trial. This analysis was applied to both the optogenetic inhibition (Figure S5A) and activation (Figure S6A) datasets. Across both datasets, we found that the reaching profiles for Post Ctr and Post Stim trials remained virtually identical (Figures S5 and S6). In the inhibition dataset, all metrics, including reach probability and latency, success probability and latency, motif transition matrix, motif ethogram, and 3D kinematics, were statistically indistinguishable between trial types (Figure S5B–J). Similarly, in the activation dataset, the motor program exhibited remarkable stability; with the exception of a minor decrease in reach latency (Figure S6C), all other primary features remained unchanged (Figure S6B–J). Collectively, the lack of carry-over effects suggests that the SNr does not play a role in the reinforcement and refinement of the forelimb motor program.

### A momentary pause in SNr activity abolishes the ongoing reaching sequence, revealing its obligatory role in real-time motor execution

Having demonstrated that SNr activity is necessary for the overall selection and vigor of reaching, we next asked whether this activity is required for the continuous, moment-by-moment control of the motor sequence. To address this, we applied transient pulses of photoinhibition, either 100 ms (Figure 4A) or 12.5 ms (Figure 4I), timed to coincide with reaching motifs. All trials where a motif (from Lift through Retract) coincided with the pulse onset were included in the analysis.

**Figure 4.**
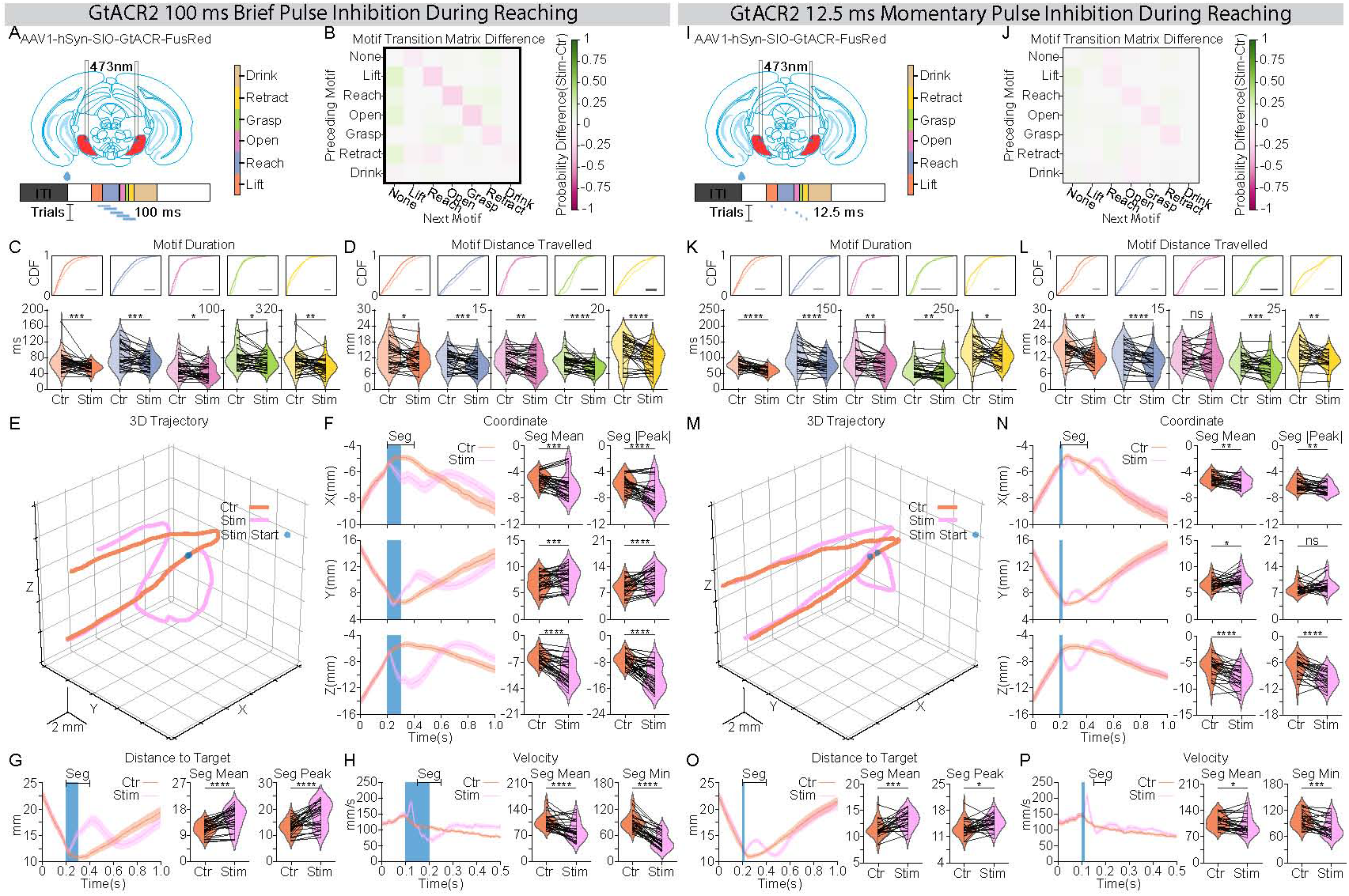
Coincident, momentary SNr inhibitory perturbation modulates online reaching kinematics. **(A)** Schematic of the experimental strategy. An inhibitory opsin (AAV1-hSyn-DIO-GtACR2-FusRed) was injected into the SNr, and an optical fiber was implanted above the region. A single 100ms photoinhibition pulse was delivered at a randomized time after water delivery onset to target reaching action sequence. The motif color map (inset) applies to all panels. **(B)** Motif transition matrix difference maps (stimulation - control). Each color block represents the difference between two conditions in transition probability from the motif on the y-axis to the motif on the x-axis. Motif sequence from a 300ms window (-40ms to 260ms aligned to laser onset) was used for quantification. The color map is shown on the right. **(C and D)** Comparison of motif duration **(C)** and motif distance travelled **(D)**. Top panels show the Cumulative Distribution Function plots of all trials; bottom panels show the violin plots of all sessions. **(E)** Average 3D forelimb trajectories (from -200ms to 800ms aligned to laser onset). The blue dot indicates laser pulse onset. Orange lines: control trials; Purple lines: stimulation trials. **(F)** Forelimb coordinates (X, Y, Z) aligned to laser pulse onset. Left: temporal traces (mean ± SEM). The blue-shaded region represents laser duration. Right: paired comparison of mean and maximum/minimum coordinates over a 200ms window starting at laser onset (Seg). **(G and H)** Forelimb distance to target **(G)** and velocity **(H)** aligned to laser pulse onset. Left: temporal traces (mean ± SEM). The blue-shaded region represents laser duration. Right: paired comparison of mean and maximum/minimum values over the designated analysis window (Seg). **(I)** as in **(A)**. But a single 12.5ms photoinhibition pulse was used instead. **(J)** as in **(B)**. **(K and L)** as in **(C and D)** respectively. **(M)** as in **(E)**. **(N)** as in **(F)**. **(O and P)** as in **(G and H)** respectively. Left panel (300 ms inhibition), n = 902 control trials and 1,319 stimulation trials from 33 sessions, 7 mice. Right panel (12.5 ms inhibition), n = 761 control trials and 911 stimulation trials from n= 26 sessions, 7 mice. Dots in the violin plots represent individual session means. Violin plots display the full data distribution density; internal box plots represent the median (center line) and the interquartile range. Wilcoxon Signed-Rank Test was used. ns no significance, *p < 0.05, **p < 0.01, ***p < 0.001, ****p < 0.0001.

Consistent with our long-term inhibition results, even a momentary pause in SNr activity was sufficient to disrupt the progression of the reaching sequence (Figure 4B-D and J-L; Video S5, 6). Motif transition analysis revealed a reduction in forward-sequence transitions along the superdiagonal, coupled with a reciprocal increase in no-reach states (leftmost column; Figure 4B, J). This behavioral disruption was further evidenced by a significant reduction in both motif duration and the distance traveled for motifs targeted by the laser pulse (Figure 4C, D and K, L). Furthermore, laser-onset-aligned 3D trajectory (Figure 4E, M), XYZ coordinate profiles (Figure 4F, N), and distance to water spout (Figure 4G, O) clearly illustrated that the reaching movement was abolished upon laser onset, with the duration of the behavioral disruption and subsequent recovery scaling with the length of the inhibitory pulse. Kinematic analysis revealed a complex velocity profile following the pause: a brief, transient increase in velocity was followed by a sustained decrease (Figure 4H, P). We reasoned that the initial velocity spike was not a direct consequence of SNr silencing, as its duration remained constant regardless of the pulse length. In contrast, the duration of the subsequent velocity reduction scaled precisely with the inhibition period, confirming that the pause of SNr activity directly reduced motor velocity in real-time. Strikingly, while the 100 ms pause induced a more profound disruption across all measured parameters (Figure 4B–H)., even the brief 12.5 ms inhibition was sufficient to abolish the ongoing motor program (Figure 4J–L).

To investigate whether this moment-by-moment control exhibits motif-specific sensitivity, trials were grouped post-hoc based on the specific action motif (Lift, Reach, Open, Grasp, or Retract) coincident with the 12.5 ms pulse onset (Figure S7A). We found that the requirement for SNr drive was non-uniform across the motor program. The ongoing action sequence was abolished during the outbound phases (Lift, Reach, and Open), followed by a re-initiation of the reach upon recovery, while the inbound phases (Grasp and Retract) were notably less affected in their sequential progression (Figure S7B). Correspondingly, laser-onset-aligned 3D trajectories (Figure S7C), coordinate profiles (Figure S7D), and distance-to-target metrics (Figure S7E) revealed a profound vulnerability from the Lift through the Grasp motifs, whereas the Retract motif remained relatively more resilient to the transient pause in SNr activity.

These findings establish that the SNr provides a continuous, motif-specific drive essential for the moment-by-moment execution of skilled reaching. Our results demonstrate that basal ganglia output is not merely a transient trigger for movement initiation or a tonic modulator of baseline vigor; rather, it is required to maintain the structural integrity and kinematic flow of the motor sequence in real-time.

### Transient SNr bursts alter the kinematics of ongoing reaching in real-time

While we have established that SNr is required for the moment-by-moment execution of skilled forelimb reaching, it is unknown if this signal serves an instructive role in real-time. To probe it, we applied transient pulses of photoactivation, either 300 ms (Figure 5A) or 12.5 ms (Figure 5I), timed to coincide with reaching motifs. All trials where a motif (from Lift through Retract) coincided with the pulse onset were included in the analysis.

**Figure 5.**
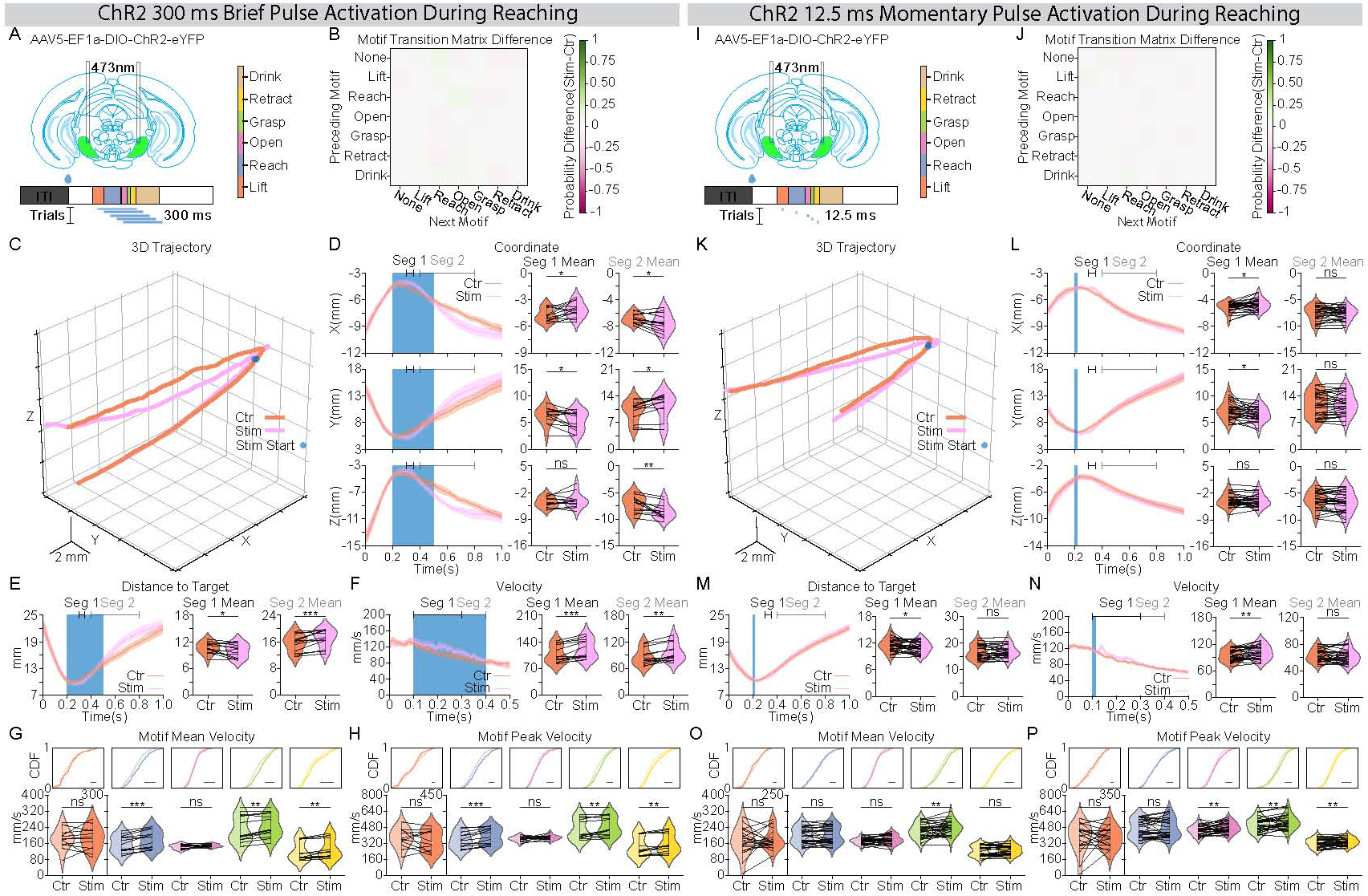
Coincident, momentary SNr excitatory perturbation modulates online reaching kinematics. **(A)** Schematic of the experimental strategy. An excitatory opsin (AAV5-EF1a-DIO-ChR2-eYFP) was injected into the SNr, and an optical fiber was implanted above the region. A single 300ms photoactivation pulse was delivered at a randomized time after water delivery onset to target reaching action sequence. The motif color map (inset) applies to all panels. **(B)** Motif transition matrix difference maps (stimulation - control). Each color block represents the difference between two conditions in transition probability from the motif on the y-axis to the motif on the x-axis. Motif sequence from a 300ms window (-40ms to 260ms aligned to laser onset) was used for quantification. The color map is shown on the right. **(C)** Average 3D forelimb trajectories (from -200ms to 800ms aligned to laser onset). The blue dot indicates laser pulse onset. Orange lines: control trials; Purple lines: stimulation trials. **(D)** Forelimb coordinates (X, Y, Z) aligned to laser pulse onset. Left: temporal traces (mean ± SEM). The blue-shaded region represents laser duration. Right: paired comparison of mean coordinates averaged over designated analysis windows (Seg 1: 300ms to 350ms; Seg 2:400ms to 800ms). **(E and F)** Forelimb distance to target **(E)** and velocity **(F)** aligned to laser pulse onset. Left: temporal traces (mean ± SEM). The blue-shaded region represents laser duration. Right: paired comparison of mean values over the designated window (distance to target: Seg 1 (300ms to 350ms), Seg 2 (400ms to 800ms); velocity: Seg 1(100ms to 300ms), Seg 2(300ms to 400ms). **(G and H)** Comparison of motif Mean Velocity **(G)** and Peak Velocity **(H)**. Top panels: Cumulative Distribution Function plots of all trials; bottom panels: violin plots of all sessions. **(I)** as in **(A)**. But a single 12.5ms photoactivation pulse was used instead. **(J)** as in **(B)**. **(K)** as in **(C)**. **(L)** as in **(D)**. **(M and N)** as in **(E and F)** respectively. **(O and P)** as in **(G and H)** respectively. Left panel (300 ms excitation), n = 493 control trials and 483 stimulation trials from 13 sessions, 4 mice. Right panel (12.5 ms excitation), n = 1,213 control trials and 1,193 stimulation trials from n= 30 sessions, 4 mice. Dots in the violin plots represent individual session means. Violin plots display the full data distribution density; internal box plots represent the median (center line) and the interquartile range. Wilcoxon Signed-Rank Test was used. ns no significance, *p < 0.05, **p < 0.01, ***p < 0.001, ****p < 0.0001.

Consistent with our long-term activation results, we found that a momentary burst in SNr activity did not impair the structural progression of the reaching sequence (Figure 5B, J and Video 5, 6). However, laser-onset-aligned 3D trajectory (Figure 5C, K), XYZ coordinate profiles (Figure 5D, L), and distance to water spout (Figure 5E, M) illustrated that the reaching movement was significantly altered upon laser onset. The altered kinematics was characterized by a distinct spatial protrusion in the trajectory (Video 5, 6), with the duration of the altered kinematics scaling with the length of the excitatory burst (Figure 5C-E and K-M). Another striking evidence of this real-time control was observed in the velocity profiles. While the 300 ms burst induced a substantially longer period of velocity enhancement, the momentary 12.5 ms burst, a duration representing only a fraction of a single motor motif, was nonetheless sufficient to drive an immediate and significant increase in forelimb velocity (Figure 5F, N). Detailed quantification across motifs confirmed that these excitatory bursts significantly enhanced both mean and peak forelimb velocities for most motifs expect Lift (Figure 5G, H and O, P).

Similarly, we grouped trials post-hoc based on the specific action motif (Lift, Reach, Open, Grasp, or Retract) coincident with the 12.5 ms pulse onset (Figure S8A). In contrast to the behavioral abolishment effect observed during SNr transient inhibition (Figure S7), the structural progression of the action sequence remained largely uninterrupted during brief activation bursts (Figure S8B). However, laser-onset-aligned 3D trajectories (Figure S8C), coordinate profiles (Figure S8D), and velocity metrics (Figure S8E) revealed an altered reaching kinematics from the Reach through the Retract motifs, whereas the Lift showed no significant deviation in its spatial trajectory or velocity profile (Figure S8C–E).

Collectively, these results reveal that rather than disrupting the progression of the forelimb sequence, a momentary burst of SNr activity preserves the structural integrity of the reach while reshaping its detailed kinematics. This demonstrates that SNr output provides a high-resolution, instructive signal that modulates the spatial and velocity profiles of reaching behavior on a moment-by-moment basis.

### Endogenous SNr population dynamics continuously represent real-time 3D kinematics

Having established that SNr activity is necessary and sufficient to drive 3D kinematics, we next investigated how endogenous SNr population dynamics encode these precise, moment-by-moment movements. To achieve this, we expressed the genetically encoded calcium indicator jGCaMP7f in the SNr and performed in vivo miniscope imaging to monitor calcium dynamics in GABAergic projection neurons at single-cell resolution during the reaching behavior (Figure 6A). Consistent with our fiber photometry results (Figure 1F-H), the population-averaged calcium signal revealed a robust increase in peri-reach activity (Figure 6B). At the single-cell level, we observed a heterogeneous mix of movement-aligned responses (Figure 6C). While movement-inhibited neurons slightly outnumbered movement-excited neurons during the initial “Lift” phase, this balance shifted as the movement progressed, with movement-excited neurons dominating the subsequent reach, grasp, retract, and drink motifs (Figure 6D). To determine how individual neurons, modulate distinct behavioral motifs, we sorted the population based on modulation magnitude during the Retract phase. We observed that population activity evolved progressively and smoothly throughout the forelimb movement sequence (from Lift to Retract). In contrast, we noted a sudden, distinct shift in population dynamics when transitioning into the orofacial “Drink” motif, suggesting distinct encoding strategies for forelimb versus consummatory movements (Figure 6E). To quantify the kinematic information embedded within this network, we trained linear regression models to predict real-time 3D kinematics using the single-cell population activity. Strikingly, the population activity robustly and continuously predicted real-time 3D trajectories (X, Y, and Z coordinates) as well as movement velocity (Figure 6F, G). Furthermore, individual neurons exhibited distributed and diverse functional weights across these different kinematic measurements (Figure 6H). Taken together, these data demonstrate that intrinsic in vivo SNr population activity contains rich, predictive representations of movement, acting as a continuous controller for moment-by-moment 3D kinematics.

**Figure 6.**
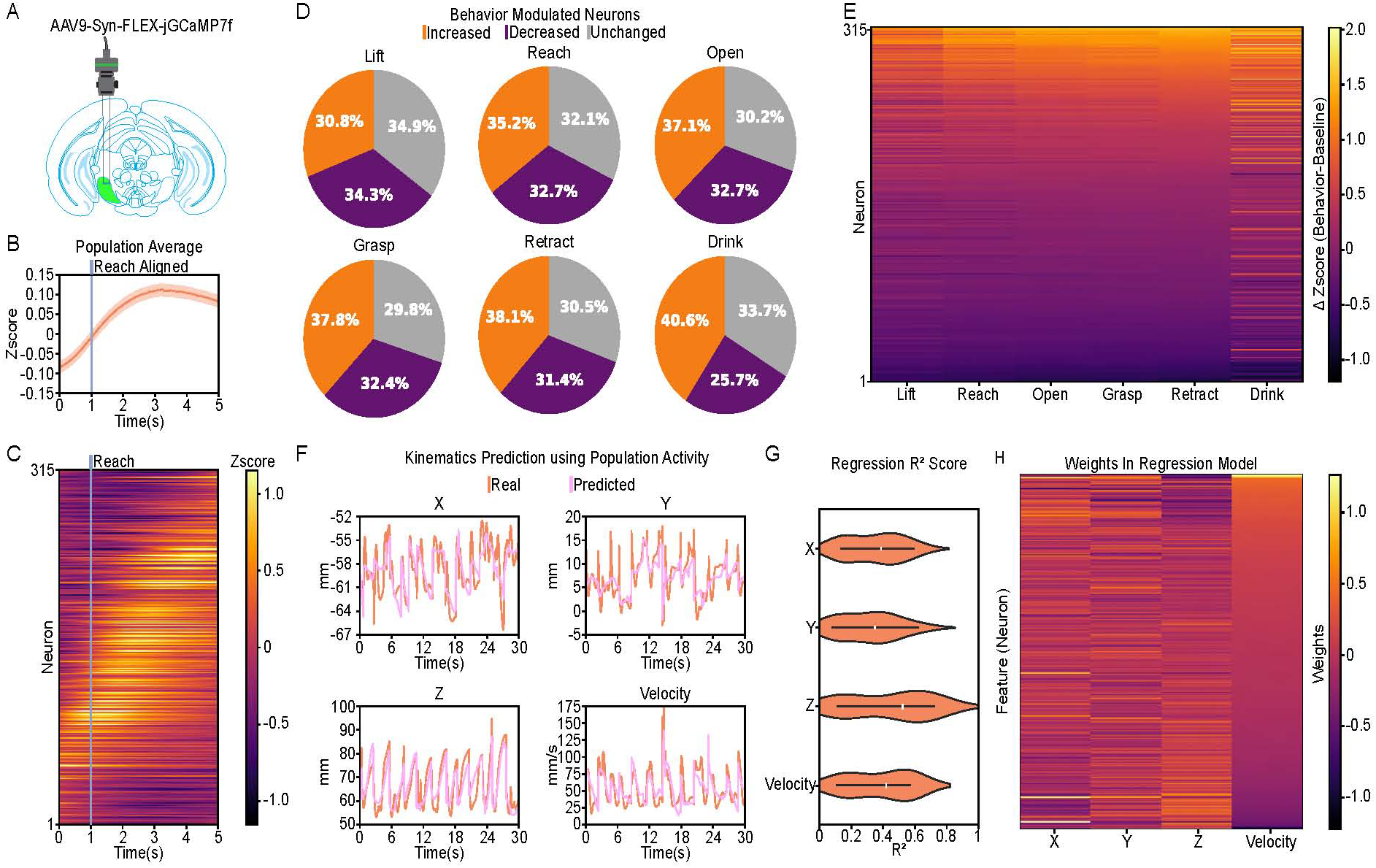
The SNr population acts as a continuous controller for moment-by-moment 3D kinematics during forelimb reaching. **(A)** Schematic of the miniscope recording strategy. AAV9-Syn-FLEX-jGCaMP7f was injected into the SNr to monitor calcium dynamics in GABAergic projection neurons at single-cell resolution during behavior. **(B)** Average population activity aligned to the onset of the Reach motif. The solid line represents the mean, and the shaded area represents SEM. **(C)** Heatmap of trial-averaged, Reach-aligned calcium activity for all recorded neurons. **(D)** Proportion of neurons exhibiting significantly increased (orange), decreased (purple), or unchanged (gray) activity during specific action motifs compared to baseline periods (Wilcoxon Signed-Rank Test). **(E)** Heatmap of the change in activity (ΔZscore; Behavior - Baseline) for all neurons across the six action motifs, sorted in descending order by modulation magnitude during the Retract motif. **(F)** Representative temporal traces comparing actual (orange) and predicted (light purple) forelimb spatial coordinates (X, Y, Z) and velocity. Predictions were generated using a linear regression model trained on concurrent SNr population activity. **(G)** Quantification of the linear regression model’s performance ($R^2$ score) in predicting moment-by-moment kinematics across sessions. Violin plots display the full data distribution density, with internal box plots representing the median (center marker) and the interquartile range. **(H)** Heatmap of normalized regression weights for each neuron across the four predicted kinematic features. Neurons (rows) are sorted in descending order by their assigned weight in the velocity model. n = 315 neurons from 600 trials, 10 sessions, 5 mice.

## Discussion

Our findings establish the substantia nigra pars reticulata (SNr) as a key component of a real-time, high-dimensional controller for skilled behavior. By integrating high-resolution 3D tracking with precisely timed perturbations and calcium imaging, we move beyond the categorical gating model to reveal a more dynamic role for basal ganglia output. We demonstrate that SNr dynamics do not merely permit movement initiation but actively dictate its kinematic flow. The paradoxical recruitment of this inhibitory output during reaching, coupled with its requirement for the moment-by-moment maintenance of motor trajectories, identifies the SNr as a critical instructive signal. By concurrently governing action selection, kinematic vigor, and real-time motor form, the SNr ensures the structural integrity and fluid execution of the complex action sequences that define motor expertise.

For decades the action selection model has dominated the study of basal ganglia function^27–29,31^. In this framework, the direct and indirect pathways are thought to exert opposing influences on the tonically active SNr to either permit or suppress motor output through downstream disinhibition^25,26^. While the discovery of co-activation between these pathways^48^ during movement led to a refined model^18,27,49,50^, where the direct pathway promotes desired actions and the indirect pathway suppresses competitors, recent evidence suggests a far more complex architecture. Specifically, the observation that the vast majority of SNr neurons show congruent, increased activity across disparate motor modalities^51^, reaching and licking, is difficult to reconcile with the refined action selection mode, which predicts that distinct motor behaviors result from opposite modulation of the firing activity of same neurons. Our study extends these observations by providing causal evidence that the SNr serves as more than a categorical switch. Through bidirectional optogenetic and chemogenetic manipulations, we demonstrate that the net activity of the SNr is engaged in both the selection of action and the titration of its kinematics (Figure 2, 3 and Figure S2-S4). Most importantly, by using transient optogenetics to perturb millisecond-scale dynamics, we reveal a critical role for the SNr in the continuous, online control of motor form (Figure 4, 5 and Figure S7, S8). Together, these results suggest that the SNr does not simply select and gate motor behavior, but actively orchestrates the detailed physical parameters of its moment-by-moment execution.

According to the dominant model, the SNr facilitates movement through a pause in its tonic firing, thereby disinhibiting downstream motor centers^25,26^. However, we demonstrate that this mechanism cannot fully account for action selection, because SNr population activity increases, rather than decreases, during skilled forelimb movement (Figure 1 and Figure 6). As this finding contradicts the canonical disinhibition model, we examined alternative circuit mechanisms that might reconcile our findings with the traditional framework. First, we addressed whether the observed population increase is a non-functional byproduct, masking a smaller, inhibited subset of neurons that actually drives the behavior. Second, we considered whether SNr activation might locally inhibit neighboring projection neurons via inhibitory axonal collaterals^44^, thereby inducing net downstream disinhibition. By using chemogenetics to modulate the SNr output globally and presumably minimizing local collateral competition, we observed behavioral effects comparable to our focal optogenetic results (Figure 2 and Figure 3). This suggests that the collective activity of the SNr, rather than a specific sub-population or modulation by axonal collaterals, plays a key role in shaping the skilled forelimb movement. In addition, we investigated whether this effect arose from post-stimulation compensation, specifically post-inhibitory rebound or post-excitatory depression, as suggested by previous work^52^. To address this, we used both chemogenetics (Figure 2 and Figure 3) and transient optogenetic perturbations (Figures 4 and Figure 5) to minimize compensatory SNr network dynamics. Through these complementary approaches, we demonstrate that the observed behavioral changes are a direct consequence of SNr drive, rather than an artifact of post-stimulation compensation. Finally, it remains possible that SNr neurons target inhibitory interneurons in downstream nuclei, providing an additional layer of nested disinhibition^53^. While our functional data cannot rule out this possibility, future studies employing cell-type-specific tracing and recording in downstream targets will be essential to map these functional connections.

Despite the long-standing popularity of the gating model, direct in vivo evidence for functional disinhibition has remained surprisingly elusive. While studies have identified movement-related firing decreases in the SNr and corresponding firing increases in its postsynaptic targets^25,26^, recent physiological surveys have found only a limited number of putative monosynaptically connected SNr-target pairs that exhibit clear inhibitory effects^16^. More tellingly, the latencies of movement-related activity changes in the SNr and its postsynaptic targets are often indistinguishable^16^. This lack of temporal offset is difficult to reconcile with a sequential disinhibition model, which would necessitate a clear delay between the release of the inhibition and the onset of downstream activity. Moreover, recent evidence suggests that the SNr has only a modulatory influence in vivo^54^, rather than showing decisive disinhibition as posited by the traditional model.

One way to reconcile the paradoxical finding that SNr excitation increases vigor, whereas even a 12.5 ms pause can abort an ongoing reach, is to interpret its output in dynamical rather than purely level-based terms. The striatonigral pathway has been proposed to transform transient, velocity-related striatal signals into sustained motor commands from the SNr output^19,33^. In such integrator-deviations from baseline decay following drive. Tonic SNr firing sets an operating point, whereas intrinsic leak reflects network stability, potentially shaped by recurrent inhibition within SNr and its connections with the rest of the BG. Our observation that global excitation accelerates reaching without impairing action selection (Figure 3, 5 and Figure S4), whereas inhibition collapses velocity and truncates motif progression (Figure 2, 4 and Figure S2, S3), is consistent with the possibility that SNr manipulations alter this dynamical stability. Global SNr excitation may shift the balance of channel-specific competition and recurrent inhibitory feedback, effectively reducing leak. In an integrator, reduced leak would allow inputs to accumulate more quickly, increasing movement velocity despite elevated absolute inhibition. Conversely, brief pauses would transiently destabilize the accumulated motor command, causing it to decay toward baseline and resulting in rapid termination of the ongoing movement. The known intranigral inhibitory microcircuit, which has been proposed to implement forms of feedback gain^44^, provides a plausible substrate for such stability modulation. Although this account remains speculative at present, the traditional model of BG fails to account for our observations. Any comprehensive model of basal ganglia function must therefore account for the critical role of SNr activity, as it dynamically stabilizes, accelerates, or terminates ongoing motor commands on a millisecond timescale.

## Methods

### Mice

All experimental procedures were conducted in accordance with the guidelines and with the approval of the Institutional Animal Care and Use Committee (IACUC) at Duke University (Protocol # 162-22-09). Adult mice (3–10 months of age) were maintained on a 12-hour light/dark cycle with ad libitum access to food and water until the start of behavioral training. Following surgical procedures, mice were singly housed and provided with environmental enrichment. This study utilized both male and female VGAT-Cre and PV-Cre transgenic lines. For behavioral experiments, mice were placed under a water restriction protocol, with their body weight monitored daily and maintained at 85–90% of their baseline weight.

### Stereotaxic injection and surgical procedures

Mice were initially anesthetized in an induction chamber with 2.0–3.0% isoflurane and subsequently secured in a stereotaxic frame (David Kopf Instruments, Tujunga, CA). Subjects received intraperitoneal injections of meloxicam (2 mg/kg) and local administration of bupivacaine for analgesia, with anesthesia maintained at 1.0–1.5% isoflurane throughout the procedure. Unilateral or bilateral craniotomies were drilled above the substantia nigra pars reticulata (SNr) at AP -3.7 mm and ML ±1.5 mm relative to bregma. Viral vectors were infused via a microinjector (Nanoject 3000, Drummond Scientific) at a rate of 1 nL/s at DV -4.4 mm relative to bregma. Following injection, the pipette remained at the target coordinate for 10 minutes to ensure adequate viral diffusion and absorption. For chemogenetic experiments, 50 nL of AAV2-hSyn-DIO-hM4D(Gi)-mCherry (Addgene #44362) or AAV2-hSyn-DIO-hM3D(Gq)-mCherry (Addgene #44361) was infused bilaterally into the SNr of VGAT-Cre or PV-Cre mice. For optogenetic experiments, 100 nL of AAV1-hSyn1-SIO-stGtACR2-FusionRed (Addgene #105677) or AAV5-EF1a-DIO-hChR2(E123T/T159C)-EYFP (Addgene #35509) was injected bilaterally into the SNr of VGAT-Cre or PV-Cre mice. Two custom-made optic fibers (105 μm core diameter, >85% transmittance, 5 mm length below the ferrule) were then implanted above the SNr at AP -3.7 mm, ML ±1.5 mm, and DV -4.4 mm relative to bregma. For fiber photometry, 100 nL of AAV9-Syn-FLEX-jGCaMP7f (Addgene #104488) was infused bilaterally into the SNr, followed by the implantation of an optic fiber (400 μm core diameter, 5 mm length below the ferrule; RWD) using the same coordinates as the optogenetic protocols. Fibers were secured to the skull using dental acrylic. For miniscope calcium imaging, 100 nL of AAV9-Syn-FLEX-jGCaMP7f (Addgene #104488) was injected unilaterally into the left SNr. A GRIN lens (6 mm x 0.5 mm; Inscopix) was implanted above the left SNr (AP -3.7 mm, ML -1.5 mm, DV -4.4 mm) and secured with dental acrylic. A baseplate for miniscope attachment was affixed to the GRIN lens 2–3 weeks post-surgery following verification of viral expression. All mice were allowed to recover for a minimum of three weeks before the commencement of behavioral experiments.

### Histology

Following the conclusion of all behavioral experiments, mice were sacrificed to confirm viral expression and verify the precise anatomical placement of optic fibers and GRIN lenses. Mice were anesthetized with 3.0% isoflurane until immobile and transcardially perfused with 0.1 M phosphate-buffered saline (PBS), followed by 4% paraformaldehyde (PFA) in PBS. Brains were carefully dissected and post-fixed in 4% PFA at 4 °C. After 24 hours, the tissue was transferred to a 30% sucrose solution for cryoprotection. Coronal sections (60 μm) were obtained using a cryostat (Leica CM 1850). Brain slices were subsequently mounted and coverslipped using Fluoroshield with DAPI (Abcam #Ab104139) to visualize cell nuclei. Fluorescent images were acquired using a ZEISS microscope (Axio Zoom.V16, RRID:SCR_016980).

### Water-reaching task

Water-restricted mice were trained in a custom-built, transparent acrylic chamber (10 cm L × 10 cm W × 15 cm H) featuring a vertical slit (8.5 mm W × 40 mm H) on the front wall. The dimensions of the slit were designed to permit reaching movements using only the right forelimb while preventing direct access to the waterspout for licking. Outside the chamber, a blunted 21G needle was used as a waterspout to deliver ∼5 μL water droplets via a solenoid valve (NResearch). Each trial was initiated with an auditory cue, and the spatial position of the spout was fixed for each subject using a three-axis manual micromanipulator. Mice were provided a 7 s response window to reach before the spout was retracted, with inter-trial intervals (ITI) randomized between 2 and 15 s. To capture the kinetics of the forelimb reaching sequence, two high-speed cameras (DMK 33UX287; The Imaging Source) were positioned perpendicularly and synchronized to record behavior at 500 Hz. An infrared (IR) light source was mounted near the cameras to ensure high-contrast illumination for kinematic tracking.

### Behavioral classification and kinematics quantification

To achieve high-resolution tracking of forelimb kinematics, two deep neural networks were trained independently using DeepLabCut (https://github.com/DeepLabCut) to digitize specific anatomical landmarks (digits 2–4, paw, mouth, and nose) across both camera views. Three-dimensional (3D) coordinates were subsequently reconstructed using the DeepLabCut 3D support pipeline following camera calibration with a checkerboard pattern^55^. These reconstructed coordinates were utilized to compute pairwise inter-bodypart coordinates and were further augmented with derived kinematic features, including instantaneous velocity and acceleration. The centroid of the three digits and the paw was used to represent the 3D coordinates of forelimb. To facilitate unbiased, high-throughput behavioral analysis, we implemented a Gated Recurrent Unit (GRU)-based classifier. This model utilized the augmented feature sets (coordinates, velocity, and acceleration) as inputs to automatically segment reaching sequences into seven discrete motor motifs: Lift, Reach, Open, Grasp, Open, Retract, and Drink. The GRU network consisted of a single hidden layer with 256 units, and was trained using a cross-entropy loss function on a large-scale, manually annotated dataset comprising 1,237,503 frames. Each identified motif is characterized by a unique and reproducible kinematic profile (Figure S1). To evaluate motor performance, motif-specific metrics were first calculated for each occurrence within a trial and then averaged to produce a single representative value per trial. The following kinematic and temporal parameters were defined. Motif number: the total count of motif occurrence per trial; Motif latency: the time elapsed from the water delivery to the first occurrence of a specific motif; Motif duration: the time spent in a single occurrence of a specific motif; Motif total duration: the cumulative time spent in a specific motif per trial; Motif distance travelled: the cumulative length of 3d trajectory traveled by the forelimb during a specific motif; Motif mean/peak velocity: the average or maximum velocity of the forelimb during a specific motif. We further defined trial-wide metrics to quantify global behavior. Latency to reach: the time from water delivery to the first Lift-Reach sequence; Latency to success: time from first Lift-Reach sequence to the first Drink motif with a duration > 600 ms. Every single parameter from each trial was then averaged to get a single numerical quantification for that session. Session-level quantifications included: Reach probability (percentage of trials containing at least one Lift-Reach sequence); Success probability (percentage of reaching trials resulting in a Drink motif > 600 ms); and Motif probability (percentage of trials containing at least one occurrence of a specific motif).

### Optogenetic experiments

Optogenetics were performed using a 473 nm blue laser (Shanghai Laser & Optics Century Co., Ltd.) delivered bilaterally, unless otherwise specified, during randomized trials. For control trials, we assigned pseudo-laser start and end times that matched the exact temporal distribution of stimulation trials within the same session to ensure rigorous quantification and comparison. All motif-specific metrics were analyzed across three distinct windows: Within-Laser, After-Laser, and Whole-trial as indicated in the figures. We conducted three sets of optogenetic inhibition experiments: 1) for fixed duration and varying intensity, a 4 s laser was delivered at water onset in 33% of trials at intensities of 10 mW, 5 mW, 2.5 mW, or 1.25 mW; 2) for fixed intensity and varying duration, a 5 mW laser was delivered at water onset for 1 s, 2 s, 3 s, or 4 s in 33% of trials; and 3) for momentary pulse inhibition, brief pulses of 12.5 ms or 100 ms were delivered in 60% of trials at optimized, mouse-specific latencies relative to water delivery to intercept specific behavioral motifs. We also performed two sets of optogenetic activation experiments using a 10 mw laser: 1) for fixed duration and varying frequency, a 4 s laser pulse (2 ms pulse width at 40 Hz or 20 Hz) was delivered at water onset in 25% of trials; and 2) for momentary pulse excitation, a laser of 12.5 ms (constant) or of 300 ms (40 Hz, 2 ms pulse width) were delivered in 50% of trials at calibrated latencies to target reaching motifs.

### Chemogenetic experiments

Chemogenetic manipulations were performed using Deschloroclozapine^56^ (DCZ; Hello Bio #HB9126), a potent and highly selective actuator for hM3D(Gq) and hM4D(Gi) DREADD receptors. Each experimental session began with 75 baseline trials to establish pre-treatment performance levels. To ensure a non-invasive administration and minimize stress-induced behavioral artifacts, mice were allowed to voluntarily consume 100 µL of water containing either saline (control) or DCZ (0.2 µg/g body weight). Following a 6-minute absorption period to allow for rapid ligand uptake, an additional 75 trials were recorded to assess the effects of SNr activation or inhibition on forelimb skilled movement. Control and DCZ sessions were performed on consecutive days, with the saline session serving as a within-subject control for the subsequent chemogenetic manipulation.

### Fiber photometry

Following a three-week recovery period for viral expression of jGCaMP7f, cell-type-specific calcium activity in the SNr was recorded during the water-reaching task. Fluorescent signals were acquired using a multi-channel fiber photometry system (RWD Life Science). Two light-emitting diodes (LEDs) were used: a 470 nm LED to excite the calcium-dependent jGCaMP7f signal and a 410 nm LED as an isosbestic control to account for movement artifacts and photo-bleaching. Light was delivered through a dichroic mirror and a multimode fiber-optic patch cord (400 µm core, 0.5 NA), with power at the fiber tip maintained between 40 mW to minimize phototoxicity. The emitted fluorescence was collected through the same patch cord, filtered, and detected by a femtowatt photoreceiver. The isosbestic 410 nm signal was fitted to the 470 nm signal to calculate the change in fluorescence (Δ F/F), defined as (F_recorded_ - F_fitted_/ F_fitted_). The resulting Δ F/F traces were Z-score normalized and temporally aligned to the behavioral metadata. The mean and peak calcium activity were calculated for each occurrence of the motor motifs (Lift, Reach, Open, Grasp, Open, and Retract) to evaluate the correlation between SNr population activity and kinematics of specific motor motifs.

### Miniscope Imaging

Following a three-week recovery period to ensure robust viral expression of jGCaMP7f, calcium dynamics in SNr projection neurons were recorded at single-cell resolution during the free-moving water-reaching task. Imaging was performed using a custom-built miniature fluorescence microscope paired with custom data acquisition software. To minimize stress and tethering-related behavioral artifacts, mice were extensively habituated to the behavioral arena while wearing a weight-matched dummy miniscope for several days prior to the start of recording experiments. To mitigate photobleaching of the calcium indicator and prevent potential phototoxicity, recording sessions were strictly capped at 60 trials per animal. Raw calcium imaging videos were preprocessed and analyzed using the MiniAn Python package^57^. The standard processing pipeline included motion correction to remove movement artifacts, background subtraction, and constrained non-negative matrix factorization (CNMF) to extract the spatial footprints and calculate temporal calcium traces of individual neurons. For trial-based analysis, the baseline activity for each trial was defined as the mean fluorescence signal during a predefined window (-2.0 s to -0.4 s relative to water delivery onset). To quantify the magnitude of behavioral modulation, a ΔZscore was calculated for each neuron by subtracting the baseline mean Zscore from the mean Zscore recorded during specific action motifs. The statistical significance of this neuronal modulation was evaluated using a Wilcoxon signed-rank test. Finally, to evaluate the relationship between neural activity and continuous behavior, a linear regression model was trained to predict moment-by-moment 3D kinematic variables (e.g., XYZ spatial coordinates and forelimb velocity) utilizing the concurrently recorded SNr population activity as inputs.

### Quantification and statistical analyses

All data are presented as mean ± SEM unless otherwise specified. Computational quantification and statistical analyses were performed using custom scripts developed in Python. Due to the non-normal distribution of behavioral and neural datasets, non-parametric statistical tests were employed throughout the study, unless otherwise noted. Specifically, for comparisons between two paired groups, the Wilcoxon signed-rank test was applied. Comparisons involving multiple groups or experimental conditions were evaluated using the Friedman test. Spearman’s rank correlation coefficient was used for correlation analysis. Results were considered statistically significant at p < 0.05. Comprehensive details regarding the specific statistical tests, exact p-values, and sample sizes are summarized in Supplementary Table 2.

## Resource Availability

### Lead Contact

Further information and requests for resources and reagents should be directed to and will be fulfilled by the lead contact, Henry H. Yin.

### Materials Availability

This study did not generate new unique reagents.

### Data and Code Availability

All data reported in this paper will be shared by the lead contact upon request.

All code used in this paper will be shared by the lead contact upon request.

Any additional information required to reanalyze the data reported in this paper is available from the lead contact upon request.

## Acknowledgments

This research was supported by NIH grant R01NS094754. We thank Rebecca Chung-Hui Yang and Lindsey Glickfeld for reading the manuscript and providing constructive feedback.

## Author Contributions

SR and HHY conceived the study and designed experiments. SR built setups, collected data, established data-analysis pipelines, analyzed data, and wrote the manuscript. HHY supervised the project and revised the manuscript.

## Declaration of Interests

The authors declare no competing interests.

---

**Figure S1.**
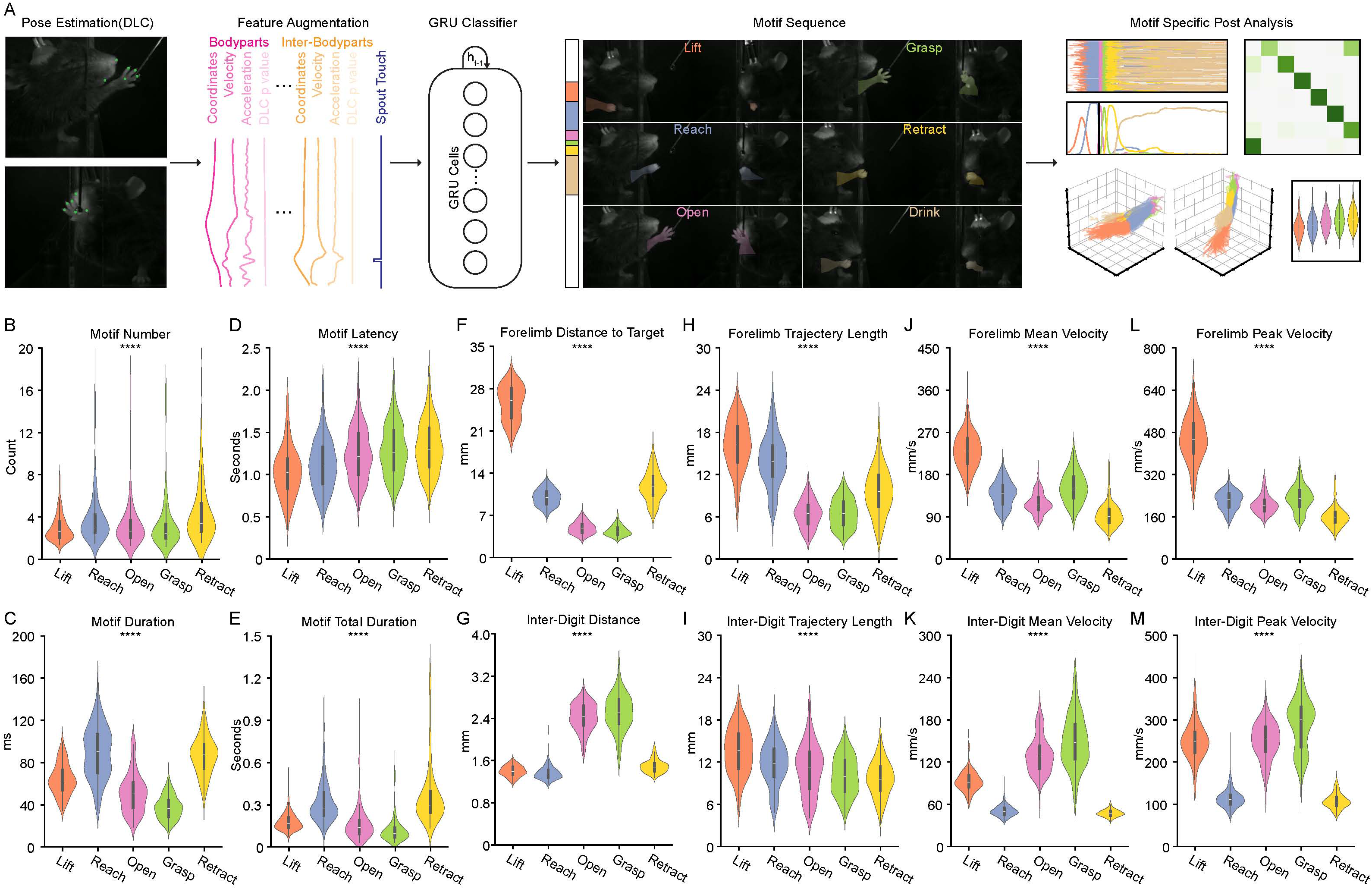
Establishment of a GRU (Gated recurrent unit)-based classifier for automated reaching sequence segmentation and motif-specific analysis, related to Figure 1. **(A)** Schematic of the behavioral analysis pipeline. DeepLabCut (DLC) was used to track six key body parts (digits 2–4, paw, mouth, and nose). Extracted coordinates were used to calculate pairwise inter-bodyparts coordinate and then were augmented with derived kinematic features, including velocity, acceleration. The augmented features were used as input to a GRU classifier for automated motif segmentation and subsequent analysis. **(B–M)** Quantification of temporal and kinematic parameters across action motifs from a large dataset (n=24498 control trials, 251 sessions from 28 mice). Violin plots display the full data distribution density; internal box plots represent the median (center line) and the interquartile range. Parameters are grouped into: **(B–E)** temporal statistics (motif count, duration, latency, and total duration); **(F–I)** coordinate related metrics (distance to target, inter-digit distance, and trajectory travelled length); and **(J–M)** velocity related metrics (mean and peak velocities for the forelimb and inter-digit spread). All quantified parameters exhibited significant variation across motifs (Friedman test, ∗∗∗∗p < 0.0001)

**Figure S2.**
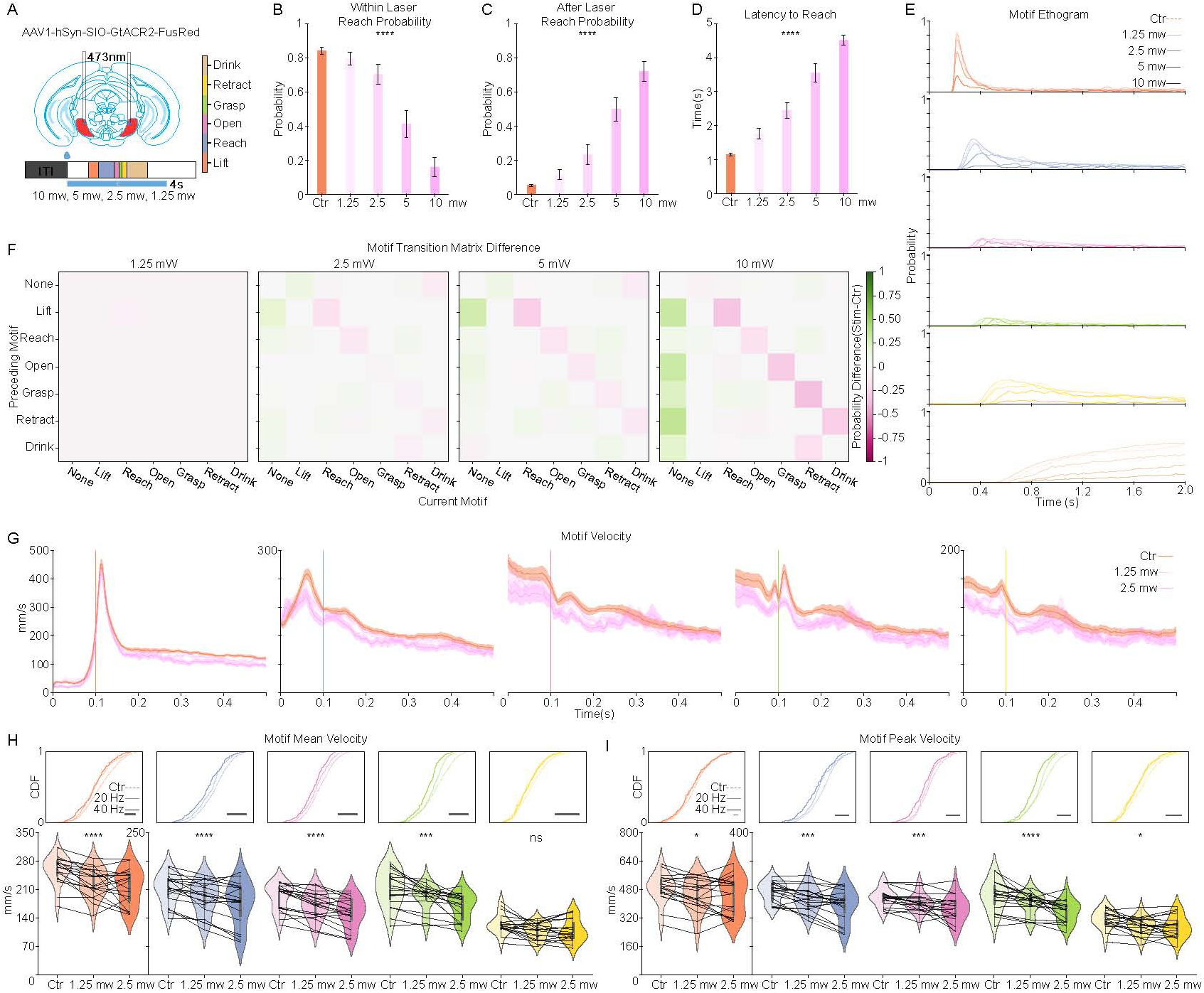
Intensity-dependent modulation of forelimb reaching behavior by SNr photoinhibition, related to Figure 2. **(A)** Schematic of the experimental strategy. An inhibitory opsin (AAV1-hSyn-DIO-GtACR2-FusRed) was injected into the SNr and an optical fiber was implanted above the region. Photoinhibition was delivered using a 473 nm blue laser at four different intensities (1.25 mW, 2.5 mW, 5 mW, 10 mW), applied randomly across trials starting at water delivery and lasting for 4 seconds. The motif color map is provided (inset). **(B–D)** Quantification of reach initiation dynamics: **(B)** Probability of reaching during laser delivery (Within Laser Reach Probability); **(C)** Probability of reaching following laser cessation (After Laser Reach Probability); and **(D)** Latency to Reach. (n = 21 sessions, 7 mice; Friedman test). **(E)** Motif ethogram aligned to the first Lift motif, stratified by inhibition intensity (n = 21 sessions, 7 mice). **(F)** Motif transition matrix difference maps, stratified by laser intensity. Each matrix displays the difference in transition probability (Inhibition - Control). n = 21 sessions, 7 mice. **(G)** Forelimb velocity profiles (mean ± SEM) aligned to the onset of Lift, Reach, Open, Grasp, and Retract motifs (left to right). Only low-intensity trials (1.25 mW and 2.5 mW) are shown, as higher intensities blocked the majority of reaching attempts (n = 19 sessions, 7 mice). **(H and I)** Quantification of motif Mean Velocity and Peak Velocity across different laser intensities. Top panels show the Cumulative Distribution Function (CDF) plots; bottom panels show the violin plots. (n = 19 sessions, 7 mice; Friedman test). Data in bar plot are shown as mean ± SEM. Dots in the bar and violin plots represent individual session means. Violin plots display the full data distribution density; internal box plots represent the median (center line) and the interquartile range. ns no significance, *p < 0.05, **p < 0.01, ***p < 0.001, ****p < 0.0001.

**Figure S3.**
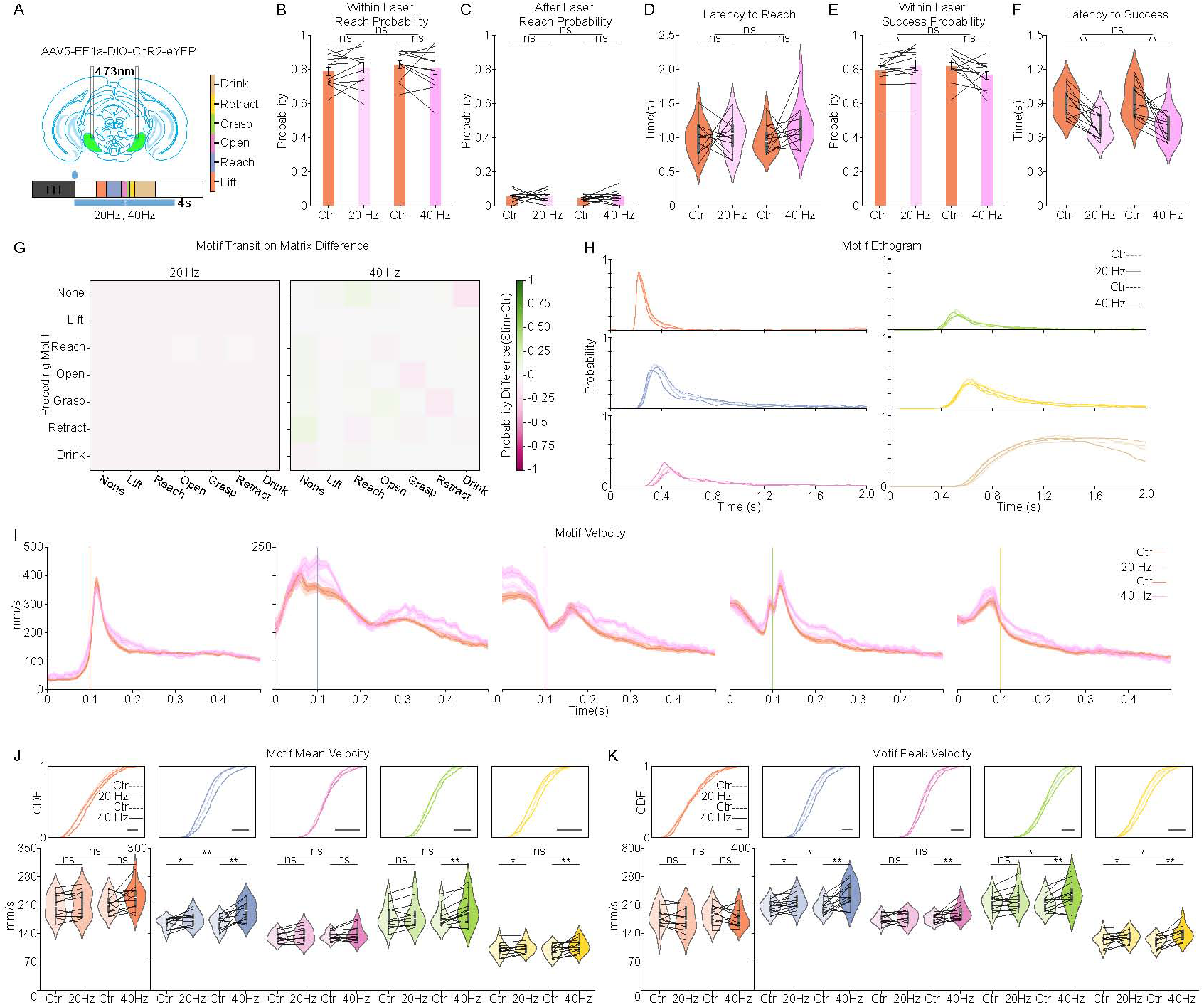
Rebound reaching following SNr inhibition exhibits enhanced kinematic signatures, related to Figure 2. **(A)** Schematic of variable-duration optogenetics inhibition. A 473 nm laser was delivered at water delivery onset for randomly interleaved durations (1 s, 2 s, 3 s, 4 s). The motif color map (inset) applies to all panels. **(B)** Motif ethograms aligned to laser onset (left) and the laser offset (right), stratified by inhibition duration in descending order. Shaded regions indicate laser-on periods (n = 808 trials, 14 sessions,5 mice). **(C)** Quantification of reach probability during (Within Laser Reach Probability) and immediately following (After Laser Reach Probability) inhibition (n = 14 sessions, 5 mice). **(D)** Cumulative Distribution Function (CDF) plots of reach latency of all trials, stratified by inhibition duration (dark to light purple) compared to controls (orange) (n = 1,652 control and 808 stimulation trials; 14 sessions, 5 mice). **(E)** Motif transition matrices. Left: reaching in control trials; right: rebound reaching in stimulation trials; middle: difference between rebound and control transition matrices. (n = 1,652 control trials and 808 stimulation trials from 14 sessions, 5 mice). **(F and G)** Comparison of reach success probability **(F)** and latency to success **(G)** between control and rebound reaches (n = 14 sessions, 4 mice; Wilcoxon Signed-Rank Test). **(H)** Superimposed motif ethograms aligned to the first Lift motif, comparing rebound (saturated colors) and control (lighter colors) reaches (n = 1,652 control and 808 stimulation trials; 14 sessions, 5 mice). **(I)** Session average forelimb velocity profiles (mean ± SEM) aligned to the onset of Lift, Reach, Open, Grasp, and Retract motifs (left to right). Orange lines: control reaches; Purple lines: rebound reaches. (n = 14 sessions, 4 mice). **(J and K)** Quantification of motif Mean Velocity **(J)** and Peak Velocity **(K)** for control vs rebound reaches. Top panels: Cumulative Distribution Function (CDF) (n = 1,652 control and 808 stimulation trials; 14 sessions, 5 mice); bottom panels: violin plots (n = 12 sessions, 4 mice; Wilcoxon Signed-Rank Test). Dots in the bar and violin plots represent individual session means. Data in bar plot are shown as mean ± SEM. Dots in the bar and violin plots represent individual session means. Violin plots display the full data distribution density; internal box plots represent the median (center line) and the interquartile range. ns no significance, *p < 0.05, **p < 0.01, ***p < 0.001, ****p < 0.0001.

**Figure S4.**
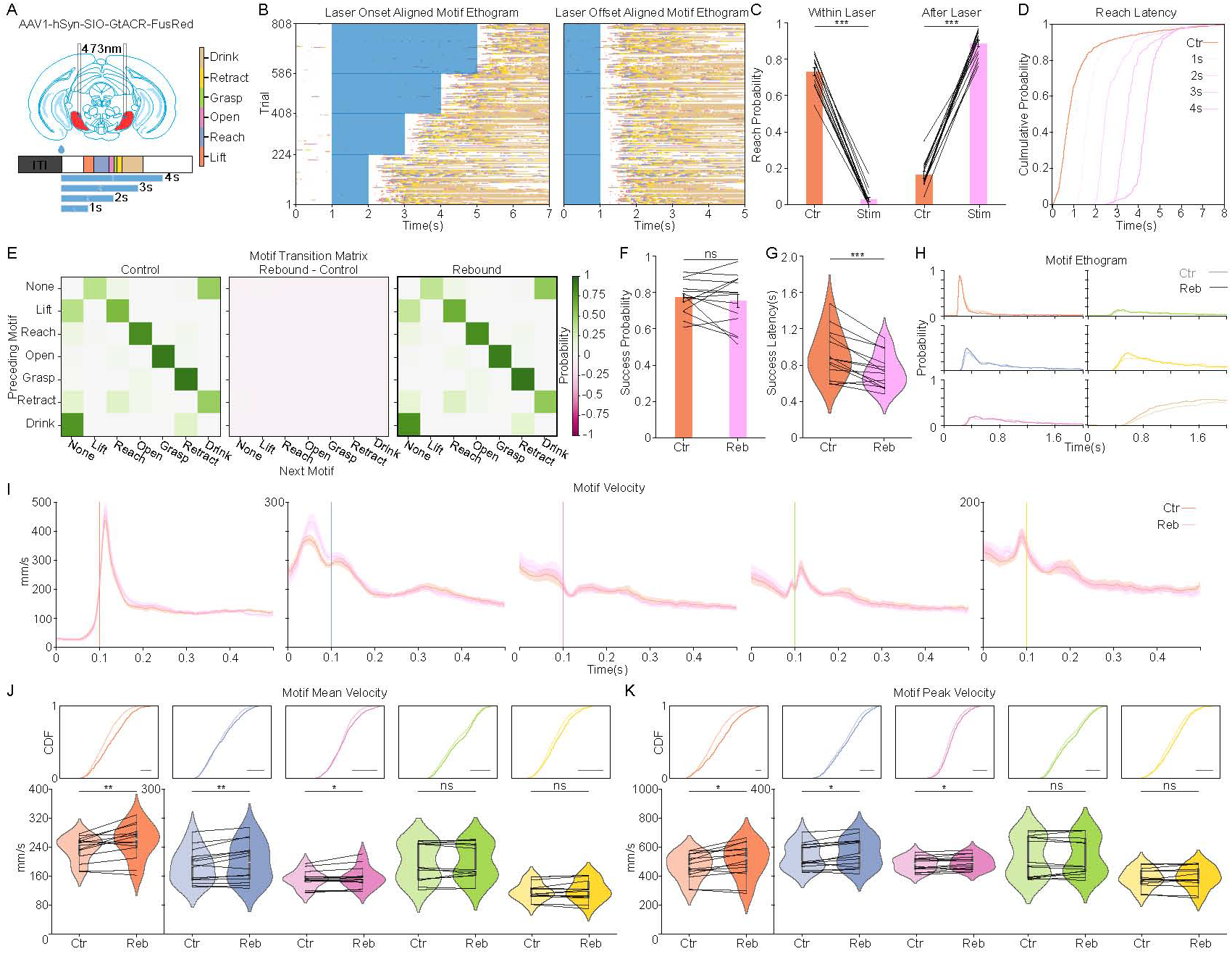
Frequency-dependent modulation of forelimb reaching behavior by SNr photoactivation, related to Figure 3. **(A)** Schematic of the optogenetic activation strategy. An excitatory opsin (AAV5-EF1a-DIO-ChR2-eYFP) was injected into the SNr and an optical fiber was implanted above the region. Photoactivation was delivered using a 473 nm blue laser at two different frequencies (20Hz, 40Hz), applied randomly across trials starting at water delivery and lasting for 4 seconds. The motif color map (inset) is used throughout the figure. **(B–D)** Quantification of reach initiation dynamics: **(B)** Probability of reaching during laser delivery (Within Laser Reach Probability); **(C)** Probability of reaching following laser cessation (After Laser Reach Probability); and **(D)** Latency to Reach. (n = 12 sessions, 4 mice; Wilcoxon Signed-Rank Test). **(E and F)** Quantification of reach success parameters: **(E)** Probability of success during laser delivery (Within Laser Success Probability); **(F)** Latency to success from reaching during laser delivery (Latency to Success). (n = 12 sessions, 4 mice; Wilcoxon Signed-Rank Test). **(G)** Motif transition matrix difference maps, stratified by laser frequency. Each matrix displays the difference in transition probability (Activation - Control). n = 12 sessions, 4 mice. **(H)** Motif ethogram aligned to the first Lift motif, stratified by inhibition intensity (n = 12 sessions, 4 mice). **(I)** Session average forelimb velocity profiles (mean ± SEM) aligned to the onset of Lift, Reach, Open, Grasp, and Retract motifs (left to right). Orange lines represent control trials, and purple lines represent activation trials. **(J and K)** Quantification of motif Mean Velocity and Peak Velocity across different laser frequencies. Top panels show the Cumulative Distribution Function (CDF) plots; bottom panels show the violin plots. (n = 12 sessions, 4 mice; Wilcoxon Signed-Rank Test). Data in bar plot are shown as mean ± SEM. Dots in the bar and violin plots represent individual session means. Violin plots display the full data distribution density; internal box plots represent the median (center line) and the interquartile range. ns no significance, *p < 0.05, **p < 0.01, ***p < 0.001, ****p < 0.0001.

**Figure S5.**
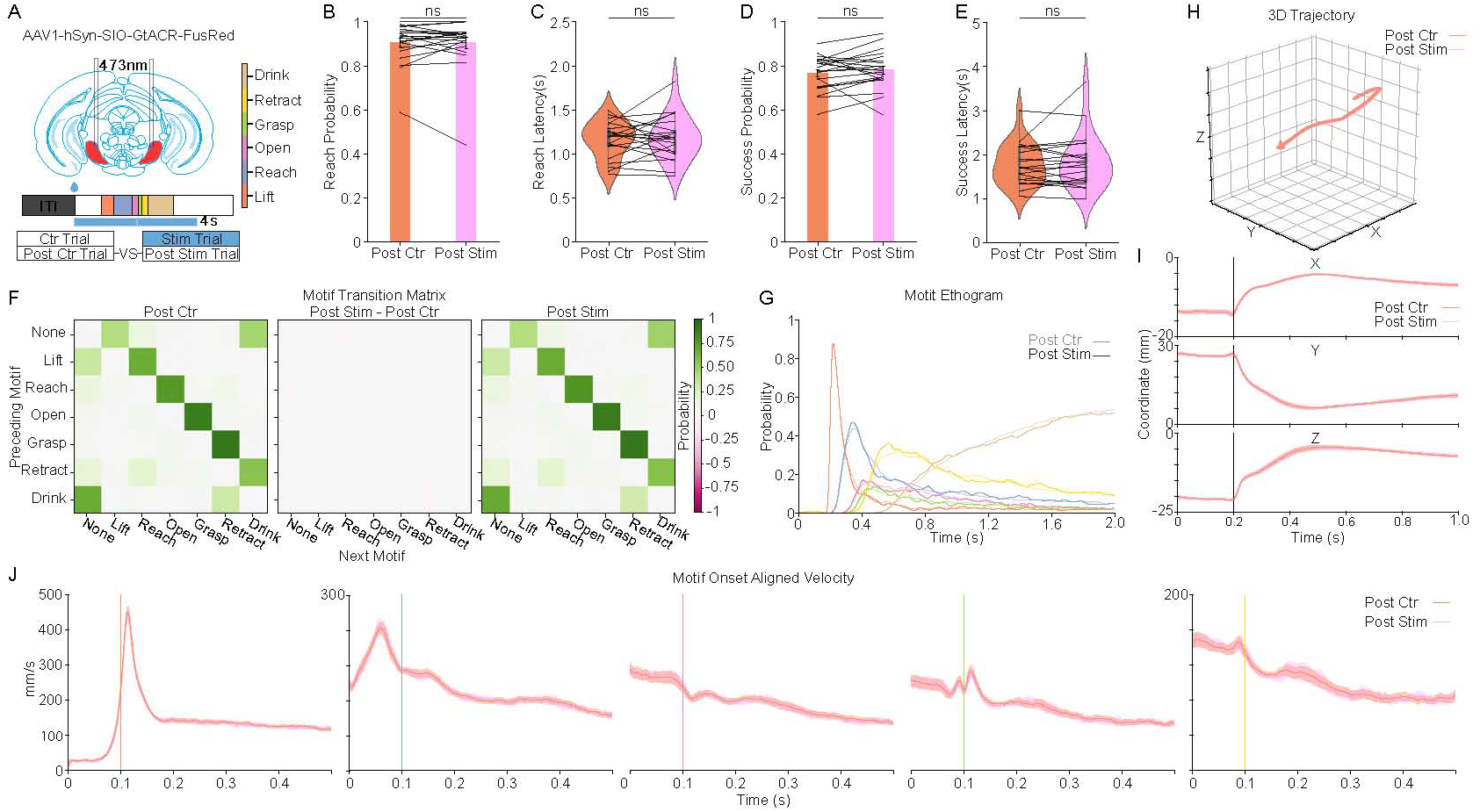
SNr photoinhibition does not produce carryover effects on subsequent control trials, related to Figure 2. **(A)** Schematic of the experimental strategy to assess carryover effects. Top: Viral expression and fiber implantation in the SNr. Middle: Diagram of the trial-based reaching task and optogenetics protocol. Bottom: Definition of trial categories; Post Ctr denotes control trials immediately following a control trial, and Post Stim denotes control trials immediately following a stimulation trial. The motif color map (inset) applies to all panels. **(B–E)** Quantification of reach dynamics comparing Post Ctr and Post Stim conditions: **(B)** Probability of reaching; **(C)** Latency to Reach; **(D)** Probability of success; and **(E)** Latency to Success. Data in bar plot are shown as mean ± SEM. Dots in the bar and violin plots represent individual session means. Violin plots display the full data distribution density; internal box plots represent the median (center line) and the interquartile range. No significant differences were observed (n = 21 sessions, 7 mice; Wilcoxon Signed-Rank Test). **(F)** Motif transition matrices for Post Ctr Trials (left) and Post Stim Trials (right). The difference matrix (middle) indicates no substantial deviation in motif sequence progression (n = 1,917 Post Ctr and 770 Post Stim Trials; 21 sessions, 7 mice). **(G)** Superimposed motif ethograms aligned to the first Lift motif, comparing Post Ctr (lighter colors) and Post Stim (saturated colors) Trials (n = 1,917 Post Ctr Trials and 770 Post Stim Trials from 21 sessions, 7 mice. **(H and I)** Forelimb reaching kinematics aligned to the first Lift motif: **(H)** Average 3D forelimb trajectories and **(I)** corresponding X, Y, Z coordinate traces. Orange lines: Post Ctr; Pink lines: Post Stim (n = 21 sessions, 7 mice). **(J)** Session average forelimb velocity profiles (mean ± SEM) aligned to the onset of Lift, Reach, Open, Grasp, and Retract motifs (left to right). Orange lines represent control trials, and purple lines represent activation trials. (n = 21 sessions, 7 mice).

**Figure S6.**
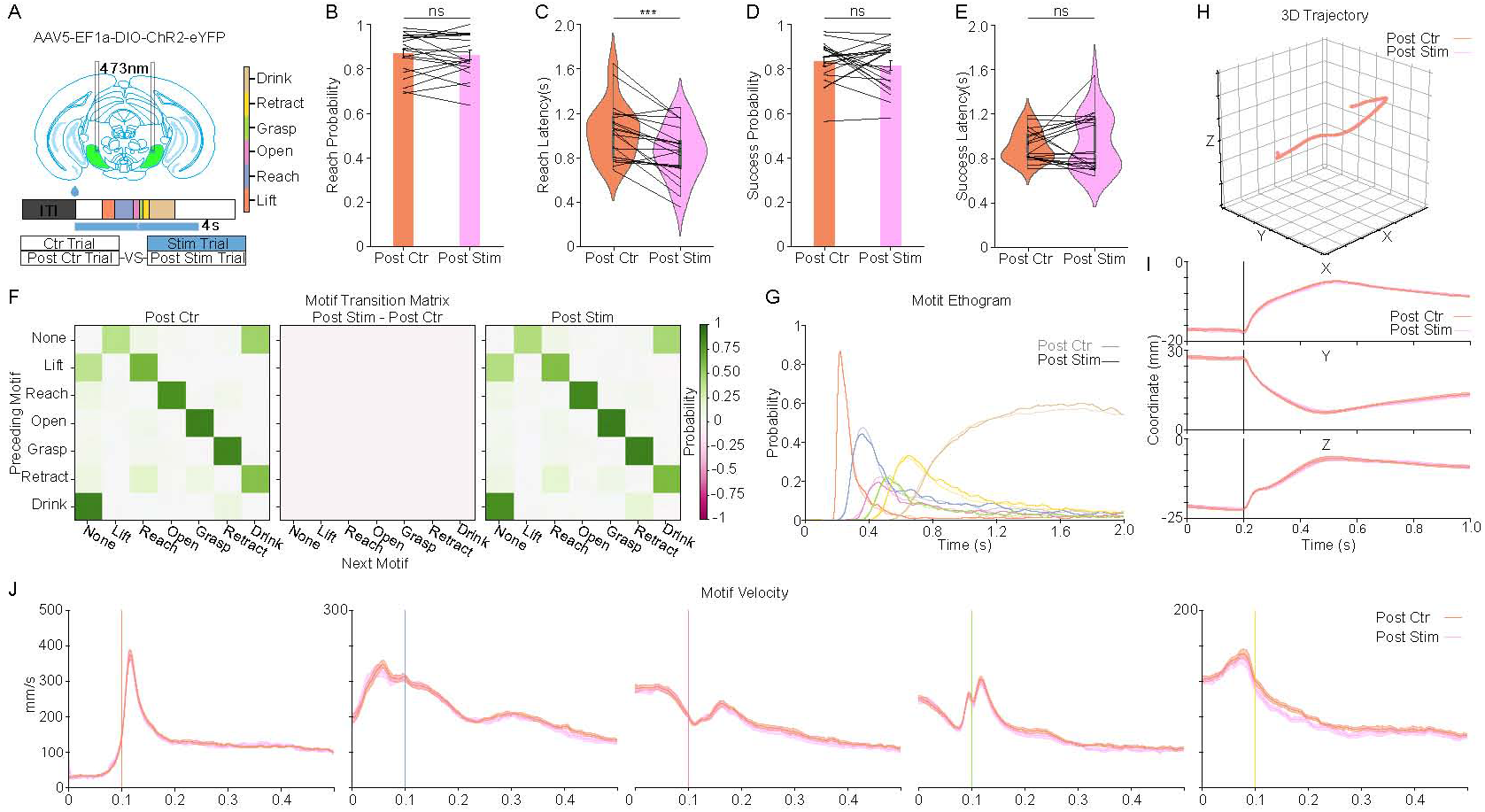
SNr photoactivation does not produce carryover effects on subsequent control trials, related to Figure 3. **(A)** Schematic of the experimental strategy to assess carryover effects. Top: Viral expression and fiber implantation in the SNr. Middle: Diagram of the trial-based reaching task and optogenetics protocol. Bottom: Definition of trial categories; Post Ctr denotes control trials immediately following a control trial, and Post Stim denotes control trials immediately following a stimulation trial. The motif color map (inset) applies to all panels. **(B–E)** Quantification of reach dynamics comparing Post Ctr and Post Stim conditions: **(B)** Probability of reaching; **(C)** Latency to Reach; **(D)** Probability of success; and **(E)** Latency to Success. Data in bar plot are shown as mean ± SEM. Dots in the bar and violin plots represent individual session means. Violin plots display the full data distribution density; internal box plots represent the median (center line) and the interquartile range. (n = 12 sessions, 4 mice; Wilcoxon Signed-Rank Test). ns no significance, *p < 0.05, **p < 0.01, ***p < 0.001, ****p < 0.0001. **(F)** Motif transition matrices for Post Ctr Trials (left) and Post Stim Trials (right). The difference matrix (middle) indicates no substantial deviation in motif sequence progression (n = 1,320 Post Ctr and 504 Post Stim Trials; 14 sessions, 4 mice). **(G)** Superimposed motif ethograms aligned to the first Lift motif, comparing Post Ctr (lighter colors) and Post Stim (saturated colors) Trials (n = 1, 320 Post Ctr Trials and 504 Post Stim Trials from 14 sessions, 4 mice. **(H and I)** Forelimb reaching kinematics aligned to the first Lift motif: **(H)** Average 3D forelimb trajectories and **(I)** corresponding X, Y, Z coordinate traces. Orange: Post Ctr; Purple: Post Stim (n = 14 sessions, 4 mice). **(J)** Session average forelimb velocity profiles (mean ± SEM) aligned to the onset of Lift, Reach, Open, Grasp, and Retract motifs (left to right). Orange lines represent control trials, and purple lines represent activation trials. (n = 14 sessions, 4 mice).

**Figure S7.**
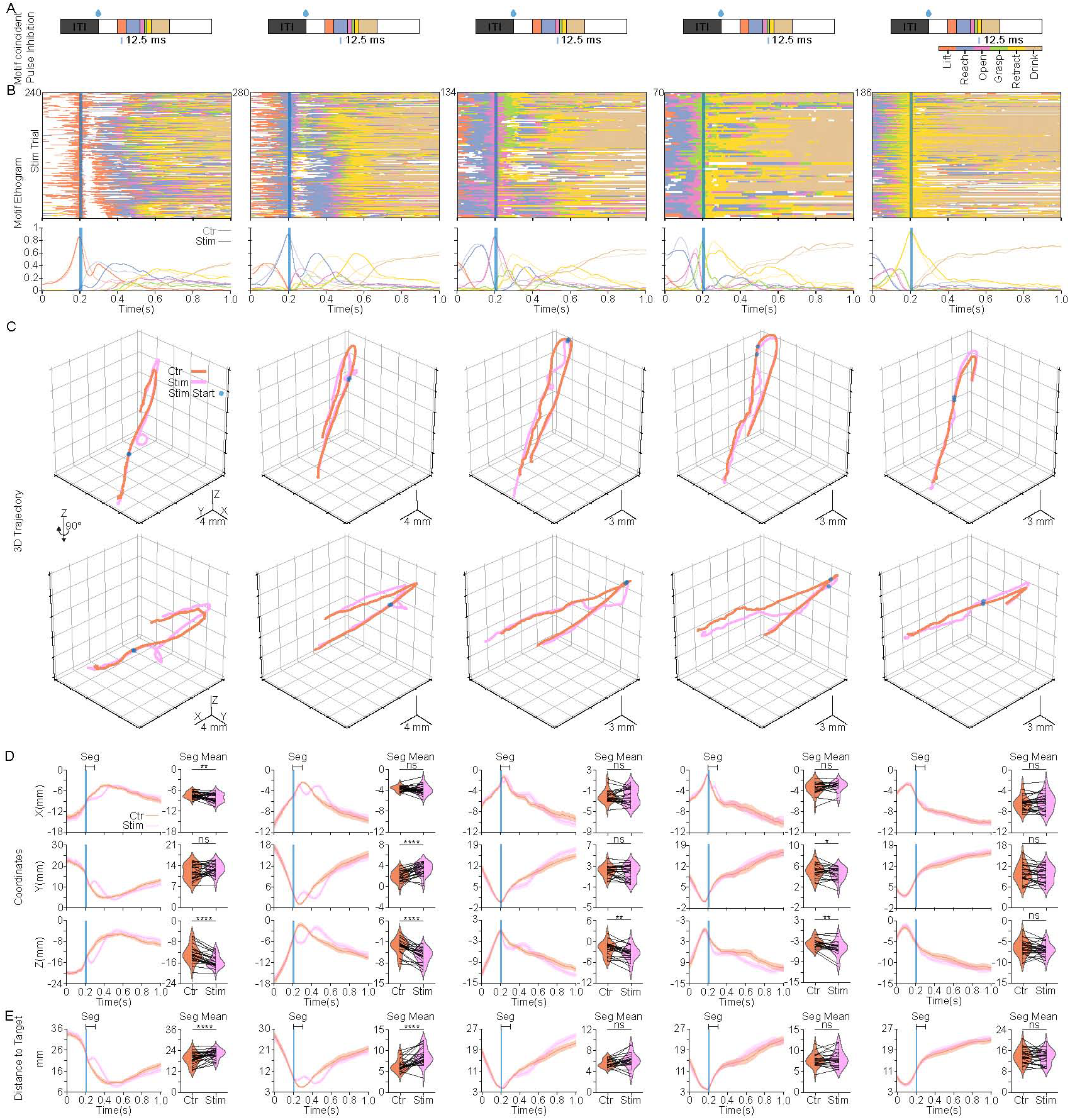
Brief SNr pulse photoinhibition produces real-time modulation of reaching kinematics, related to Figure 4. **(A)** Schematic of the experimental strategy. A single 12.5 ms photoinhibition pulse was delivered at a randomized time after water delivery onset. Trials were grouped and analyzed post-hoc based on the action motif coincident with pulse onset (Lift, Reach, Open, Grasp, or Retract). The motif color map (inset) applies to all panels. **(B)** Motif ethograms (top) and corresponding probability (bottom) aligned to laser pulse onset, stratified by the coincident motif (left to right: Lift, Reach, Open, Grasp, or Retract). The blue-shaded region indicates the laser duration. (Total n = 898 trials from 26 sessions, 7 mice; Lift: 240, Reach: 280, Open: 134, Grasp: 70, Retract: 186 trials). **(C)** Average 3D forelimb trajectories aligned to laser pulse onset (blue dots), stratified by the coincident motif. Orange lines: control trials; Purple lines: stimulation trials. (n = 26 sessions, 7 mice). **(D)** Forelimb coordinates (X, Y, Z) aligned to laser pulse onset. Left: temporal traces (mean ± SEM). Right: paired comparison of mean coordinates averaged over a 100 ms window starting at laser onset (Seg). Violin plots display the full data distribution density; internal box plots represent the median (center line) and the interquartile range. Orange: control trials; Purple: stimulation trials. (n = 26 sessions, 7 mice; Wilcoxon Signed-Rank Test). ns no significance, *p < 0.05, **p < 0.01, ***p < 0.001, ****p < 0.0001. **(E)** Forelimb distance to target aligned to laser pulse onset. Left: temporal traces (mean ± SEM). Right: paired comparison of mean distance averaged over a 100 ms window starting at laser onset (Seg). Violin plots display the full data distribution density; internal box plots represent the median (center line) and the interquartile range. Orange: control trials; Purple: stimulation trials. (n = 26 sessions, 7 mice; Wilcoxon Signed-Rank Test). ns no significance, *p < 0.05, **p < 0.01, ***p < 0.001, ****p < 0.0001.

**Figure S8.**
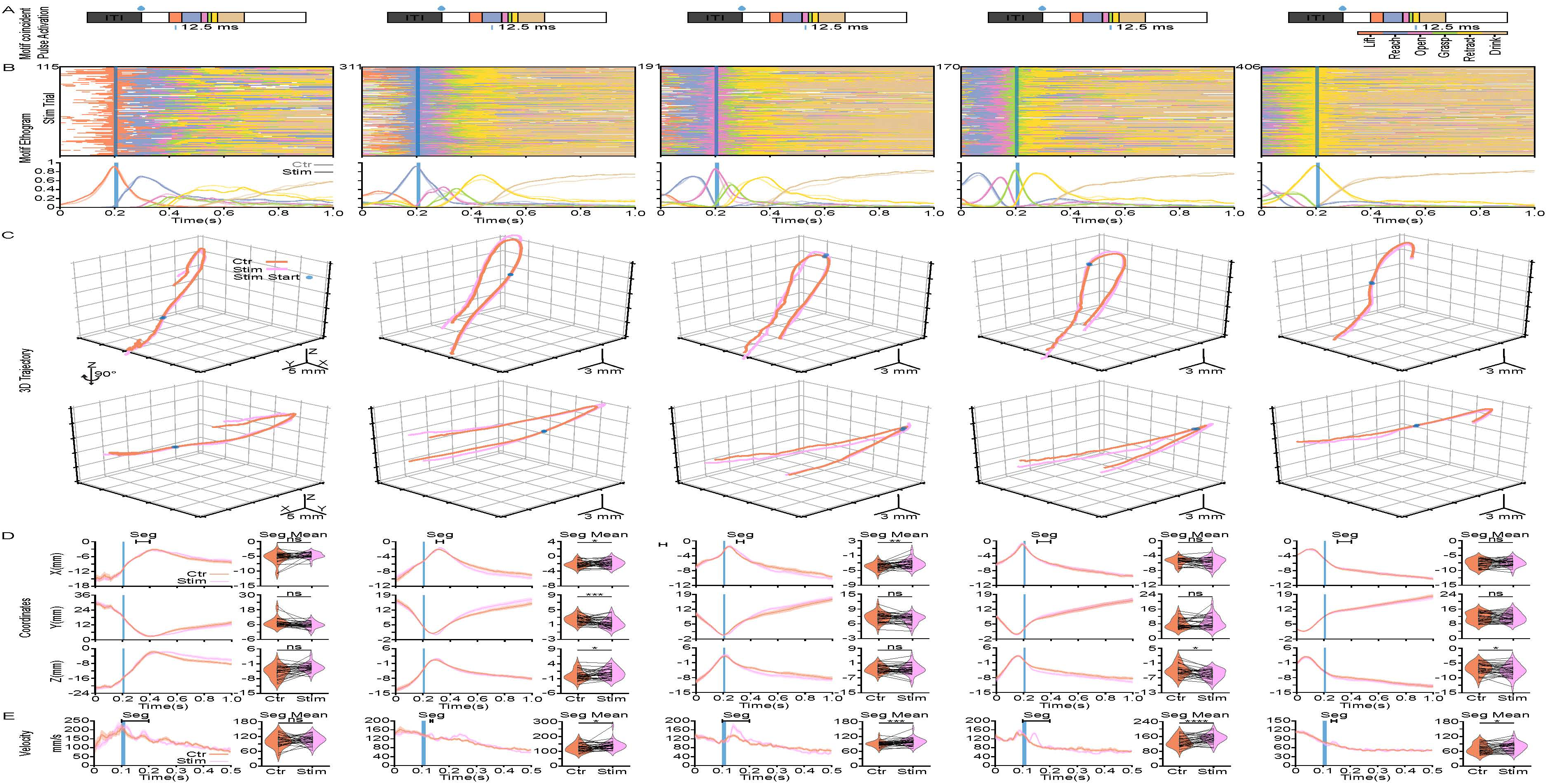
Brief SNr pulse photoactivation produces real-time modulation of reaching kinematics, related to Figure 5. **(A)** Schematic of the experimental strategy. A single 12.5 ms photoinhibition pulse was delivered at a randomized time after water delivery onset. Trials were grouped and analyzed post-hoc based on the action motif coincident with pulse onset (Lift, Reach, Open, Grasp, or Retract). The motif color map (inset) applies to all panels. **(B)** Motif ethograms (top) and corresponding probability (bottom) aligned to laser pulse onset, stratified by the coincident motif (left to right: Lift, Reach, Open, Grasp, or Retract). The blue-shaded region indicates the laser duration. (Total n = 1193 trials from 30 sessions, 4 mice; Lift: 115, Reach: 311, Open: 191, Grasp: 170, Retract: 406 trials). **(C)** Average 3D forelimb trajectories aligned to laser pulse onset (blue dots), stratified by the coincident motif. Orange lines: control trials; Purple lines: stimulation trials. (n = 30 sessions, 4 mice). **(D)** Forelimb coordinates (X, Y, Z) aligned to laser pulse onset. Left: temporal traces (mean ± SEM). Right: paired comparison of mean coordinates averaged over a defined window (Seg). Violin plots display the full data distribution density; internal box plots represent the median (center line) and the interquartile range. Orange: control trials; Purple: stimulation trials. (n = 30 sessions, 4 mice; Wilcoxon Signed-Rank Test). ns no significance, *p < 0.05, **p < 0.01, ***p < 0.001, ****p < 0.0001. **(E)** Forelimb velocity aligned to laser pulse onset. Left: temporal traces (mean ± SEM). Right: paired comparison of mean velocity averaged over a defined window (Seg). Violin plots display the full data distribution density; internal box plots represent the median (center line) and the interquartile range. Orange: control trials; Purple: stimulation trials. (n = 30 sessions, 4 mice; Wilcoxon Signed-Rank Test). ns no significance, *p < 0.05, **p < 0.01, ***p < 0.001, ****p < 0.0001.

**Supplementary Video 1. Reaching task structure and skilled forelimb motifs.** Example trial demonstrating the behavioral setup, task progression, and motor motifs in the skilled forelimb reaching task. Related to Figure 1.

**Supplementary Video 2. 3D trajectory reconstructions of forelimb reaching motifs.** Reconstructed 3D trajectories of all trials from an example session, color-coded by action motif. The video shows 500 total frames per trial, beginning 100 frames prior to the first Grasp motif following water delivery. Trajectory tails are set to a 50-frame fade. Related to Figure 1.

**Supplementary Video 3. Behavioral effects of SNr manipulation.** Representative trials showing the effects of 4-s optogenetic inhibition, chemoinhibition, 4-s optogenetic activation, and chemoactivation on forelimb skilled behavior. Related to Figures 2 and 3.

**Supplementary Video 4. Session-averaged 3D trajectories under SNr manipulation.** Reconstructed 3D trajectories (session averages) comparing experimental (purple) and control (orange) conditions. Data includes 4-s optogenetic inhibition, chemoinhibition, 4-s optogenetic activation, and chemoactivation. Each curve represents a single session. Video shows 250 frames per session, starting 50 frames prior to laser onset. Related to Figures 2 and 3.

**Supplementary Video 5. Behavioral effects of short optogenetic manipulation.** Example trials demonstrating the effects of short optogenetics manipulation coinciding with specific motor motifs: 100-ms and 12.5-ms inhibition coinciding with Lift; 100-ms and 12.5-ms inhibition coinciding with Reach; and 300-ms and 12.5-ms activation coinciding with Reach. Related to Figures 4 and 5.

**Supplementary Video 6. Reconstructed 3D trajectories under short optogenetic manipulation.** 3D trajectories of all trials from representative mice during short optogenetic manipulations. Includes 100-ms and 12.5-ms inhibition (Lift and Reach) and 300-ms and 12.5-ms activation (Lift-to-Retract). Purple: Inhibition; Orange: Control. Each curve represents one trial color-coded by action motif. Video shows 250 frames per trial, starting 50 frames prior to laser onset. Related to Figures 4 and 5.

**Supplementary Table 1.** Spearman’s rank correlation coefficient of Motif specific population calcium Zscore and forelimb movement kinematics, related to Figure 1.

|  | Ipsilateral |  | Contralateral |  | Ipsilateral |  |  |  | Contralateral |  |  |  |
| --- | --- | --- | --- | --- | --- | --- | --- | --- | --- | --- | --- | --- |
| Zscore | Mean | Peak | Mean | Peak | Mean | Peak | Mean | Peak | Mean | Peak | Mean | Peak |
|  | Motif Trajectory Length |  |  |  | Vm | Vp | Vm | Vp | Vm | Vp | Vm | Vp |
| Lift | R: 0.140<br>p: 0.544 | R: 0.110<br>p: 0.634 | R: 0.061<br>p: 0.793 | R: 0.090<br>p: 0.695 | R: -0.349<br>p: 0.121 | R: -0.238<br>p: 0.300 | R: -0.347<br>p: 0.124 | R: -0.258<br>p: 0.258 | R: -0.266<br>p: 0.243 | R: -0.230<br>p: 0.316 | R: -0.260<br>p: 0.256 | R: -0.206<br>p: 0.369 |
| Reach | R: 0.105<br>p: 0.650 | R: 0.049<br>p: 0.832 | R: 0.404<br>p: 0.069 | R: 0.518<br>p: 0.016 | R: 0.208<br>p: 0.366 | R: 0.119<br>p: 0.606 | R: 0.114<br>p: 0.622 | R: 0.057<br>p: 0.806 | R: 0.300<br>p: 0.186 | R: 0.401<br>p: 0.071 | R: 0.390<br>p: 0.081 | R: 0.505<br>p: 0.019 |
| Open | R: 0.526<br>p: 0.014 | R: 0.526<br>p: 0.014 | R: 0.560<br>p: 0.008 | R: 0.588<br>p: 0.005 | R: 0.496<br>p: 0.022 | R: 0.553<br>p: 0.009 | R: 0.479<br>p: 0.028 | R: 0.547<br>p: 0.010 | R: 0.409<br>p: 0.066 | R: 0.617<br>p: 0.003 | R: 0.436<br>p: 0.048 | R: 0.638<br>p: 0.002 |
| Grasp | R: 0.009<br>p: 0.969 | R: -0.032<br>p: 0.889 | R: 0.591<br>p: 0.005 | R: 0.604<br>p: 0.004 | R: 0.406<br>p: 0.067 | R: 0.390<br>p: 0.081 | R: 0.365<br>p: 0.104 | R: 0.348<br>p: 0.122 | R: 0.573<br>p: 0.007 | R: 0.562<br>p: 0.008 | R: 0.577<br>p: 0.006 | R: 0.562<br>p: 0.008 |
| Retract | R: 0.094<br>p: 0.687 | R: 0.103<br>p: 0.658 | R: 0.625<br>p: 0.002 | R: 0.630<br>p: 0.002 | R: 0.503<br>p: 0.020 | R: 0.291<br>p: 0.201 | R: 0.509<br>p: 0.018 | R: 0.294<br>p: 0.197 | R: 0.622<br>p: 0.003 | R: 0.504<br>p: 0.020 | R: 0.622<br>p: 0.003 | R: 0.512<br>p: 0.018 |

**Supplementary Table 2.** Statistics summary table.

| Fig | Statistic Method | Sample Size | P value |
| --- | --- | --- | --- |
| 1I | Spearman's rank correlation coefficient | 21 sessions from 7 mice | See Table 1 |
| 1J | Pearson correlation coefficient | 21 sessions from 7 mice | **Top:** R = 0.571, p = 0.007<br>**Bottom:** R = 0.658, p = 0.001 |
| 2B | Wilcoxon Signed-Rank Test | 21 sessions from 7 mice | < 0.0001 |
| 2C | Wilcoxon Signed-Rank Test | 21 sessions from 7 mice | 0.272 |
| 2E | Wilcoxon Signed-Rank Test | 21 sessions from 7 mice | **From left to right:**<br>< 0.0001 \| < 0.0001 \| < 0.0001<br>< 0.0001 \| < 0.0001 \| < 0.0001 |
| 2F | Wilcoxon Signed-Rank Test | 21 sessions from 7 mice | **From left to right:**<br>< 0.0001 \| < 0.0001 \| < 0.0001<br>< 0.0001 \| < 0.0001 \| 0.0002 |
| 2G | Wilcoxon Signed-Rank Test | 21 sessions from 7 mice | **From left to right:**<br>0.014 \| < 0.0001 \| < 0.0001<br>< 0.0001 \| < 0.0001 \| < 0.0001 |
| 2H | Wilcoxon Signed-Rank Test | 21 sessions from 7 mice | **From left to right:**<br>0.046 \| < 0.0001 \| 0.0002<br>< 0.0001 \| < 0.0001 \| < 0.0001 |
| 2J | Wilcoxon Signed-Rank Test | 21 sessions from 7 mice | **From left to right:**<br>0.0002 \| 0.0003 \| < 0.0001 \| < 0.0001 \| 0.070 |
| 2K | Wilcoxon Signed-Rank Test | 21 sessions from 7 mice | **From left to right:**<br>0.002 \| 0.0002 \| < 0.0001 \| < 0.0001 \| 0.035 |
| 2M | Wilcoxon Signed-Rank Test | 15 sessions from 5 mice | 0.389 |
| 2N | Wilcoxon Signed-Rank Test | 15 sessions from 5 mice | 0.0001 |
| 2P | Wilcoxon Signed-Rank Test | 15 sessions from 5 mice | **From left to right:**<br>0.598 \| 0.086 \| 0.124 \| 0.125 \| 0.098 \| 0.035 |
| 2Q | Wilcoxon Signed-Rank Test | 15 sessions from 5 mice | **From left to right:**<br>0.720 \| 0.359 \| 0.229 \| 0.303 \| 0.252 \| 0.022 |
| 2R | Wilcoxon Signed-Rank Test | 15 sessions from 5 mice | **From left to right:**<br>0.0004 \| 0.002 \| 0.004 \| 0.005 \| 0.002 \| 0.0001 |
| 2S | Wilcoxon Signed-Rank Test | 15 sessions from 5 mice | **From left to right:**<br>0.0001 \| 0.010 \| 0.048 \| 0.005 \| 0.012 \| 0.0004 |
| 2U | Wilcoxon Signed-Rank Test | 15 sessions from 5 mice | **From left to right:**<br>0.002 \| < 0.0001 \| 0.389 \| 0.083 \| 0.026 |
| 2V | Wilcoxon Signed-Rank Test | 15 sessions from 5 mice | **From left to right:**<br>< 0.0001 \| < 0.0001 \| 0.003 \| 0.018 \| 0.008 |
| 3B | Wilcoxon Signed-Rank Test | 21 sessions from 4 mice | 0.733 |
| 3C | Wilcoxon Signed-Rank Test | 21 sessions from 4 mice | < 0.0001 |
| 3E | Wilcoxon Signed-Rank Test | 21 sessions from 4 mice | **From left to right:**<br>0.683 \| 0.683 \| 0.683 \| 0.733 \| 0.865 \| 0.562 |
| 3F | Wilcoxon Signed-Rank Test | 21 sessions from 4 mice | **From left to right:**<br>0.393 \| 0.203 \| 0.035 \| 0.179 \| 0.320 \| 0.065 |
| 3G | Wilcoxon Signed-Rank Test | 21 sessions from 4 mice | **From left to right:**<br>< 0.0001 \| < 0.0001 \| 0.229<br>0.946 \| 0.060 \| 0.168 |
| 3H | Wilcoxon Signed-Rank Test | 21 sessions from 4 mice | **From left to right:**<br>0.006 \| 0.243 \| 0.002 \| 0.016 \| 0.355 \| 0.096 |
| 3J | Wilcoxon Signed-Rank Test | 21 sessions from 4 mice | **From left to right:**<br>0.018 \| 0.002 \| 0.119 \| 0.007 \| 0.004 |
| 3K | Wilcoxon Signed-Rank Test | 21 sessions from 4 mice | **From left to right:**<br>0.272 \| 0.0006 \| 0.006 \| 0.002 \| 0.0004 |
| 3M | Wilcoxon Signed-Rank Test | 12 sessions from 4 mice | 0.110 |
| 3N | Wilcoxon Signed-Rank Test | 12 sessions from 4 mice | 0.007 |
| 3P | Wilcoxon Signed-Rank Test | 12 sessions from 4 mice | **From left to right:**<br>0.212 \| 0.306 \| 0.350 \| 0.266 \| 0.306 \| 0.358 |
| 3Q | Wilcoxon Signed-Rank Test | 12 sessions from 4 mice | **From left to right:**<br>0.970 \| 0.622 \| 0.301 \| 0.266 \| 0.380 \| 0.092 |
| 3R | Wilcoxon Signed-Rank Test | 12 sessions from 4 mice | **From left to right:**<br>0.016 \| 0.007 \| 0.569 \| 0.622 \| 0.005 \| 0.970 |
| 3S | Wilcoxon Signed-Rank Test | 12 sessions from 4 mice | **From left to right:**<br>0.470 \| 0.850 \| 0.003 \| 0.424 \| 0.151 \| 0.470 |
| 3U | Wilcoxon Signed-Rank Test | 12 sessions from 4 mice | **From left to right:**<br>0.005 \| 0.001 \| 0.005 \| 0.042 \| 0.002 |
| 3V | Wilcoxon Signed-Rank Test | 12 sessions from 4 mice | **From left to right:**<br>0.380 \| 0.001 \| 0.0015 \| 0.027 \| 0.027 |
| 4C | Wilcoxon Signed-Rank Test | 33 sessions from 7 mice | **Left to Right (Lift to Retract):**<br>0.0002 \| 0.0001 \| 0.016 \| 0.012 \| 0.009 |
| 4D | Wilcoxon Signed-Rank Test | 33 sessions from 7 mice | **Left to Right (Lift to Retract):**<br>0.012 \| 0.0001 \| 0.005 \| < 0.0001 \| < 0.0001 |
| 4F | Wilcoxon Signed-Rank Test | 33 sessions from 7 mice | **X Seg Mean:** 0.0002<br>**X Seg \|Peak\|:** < 0.0001 |
|  |  |  | **Y Seg Mean:** 0.0004<br>**Y Seg \|Peak\|:** < 0.0001<br>**Z Seg Mean:** < 0.0001<br>**Z Seg \|Peak\|:** < 0.0001 |
| 4G | Wilcoxon Signed-Rank Test | 33 sessions from 7 mice | **Mean:** < 0.0001<br>**Peak:** < 0.0001 |
| 4H | Wilcoxon Signed-Rank Test | 33 sessions from 7 mice | **Mean:** < 0.0001<br>**Min:** < 0.0001 |
| 4K | Wilcoxon Signed-Rank Test | 26 sessions from 7 mice | **Left to Right (Lift to Retract):**<br>< 0.0001 \| < 0.0001 \| 0.002 \| 0.007 \| 0.012 |
| 4L | Wilcoxon Signed-Rank Test | 26 sessions from 7 mice | **Left to Right (Lift to Retract):**<br>0.005 \| < 0.0001 \| 0.277 \| 0.0001 \| 0.006 |
| 4N | Wilcoxon Signed-Rank Test | 26 sessions from 7 mice | **X Seg Mean:** 0.006<br>**X Seg \|Peak\|:** 0.006<br>**Y Seg Mean:** 0.025<br>**Y Seg \|Peak\|:** 0.089<br>**Z Seg Mean:** < 0.0001<br>**Z Seg \|Peak\|:** < 0.0001 |
| 4O | Wilcoxon Signed-Rank Test | 26 sessions from 7 mice | **Mean:** 0.0005<br>**Peak:** 0.012 |
| 4P | Wilcoxon Signed-Rank Test | 26 sessions from 7 mice | **Mean:** 0.014<br>**Min:** 0.0001 |
| 5D | Wilcoxon Signed-Rank Test | 13 sessions from 4 mice | **X Seg 1:** 0.048<br>**X Seg 2:** 0.017<br>**Y Seg 1:** 0.017<br>**Y Seg 2:** 0.027<br>**Z Seg 1:** 0.376<br>**Z Seg 2:** 0.001 |
| 5E | Wilcoxon Signed-Rank Test | 13 sessions from 4 mice | **Seg 1:** 0.027<br>**Seg 2:** 0.0007 |
| 5F | Wilcoxon Signed-Rank Test | 13 sessions from 4 mice | **Seg 1:** 0.0005<br>**Seg 2:** 0.001 |
| 5G | Wilcoxon Signed-Rank Test | 13 sessions from 4 mice | **Left to Right (Lift to Retract):**<br>0.970 \| 0.0002 \| 0.305 \| 0.002 \| 0.001 |
| 5H | Wilcoxon Signed-Rank Test | 13 sessions from 4 mice | **Left to Right (Lift to Retract):**<br>0.380 \| 0.0002 \| 0.216 \| 0.005 \| 0.001 |
| 5L | Wilcoxon Signed-Rank Test | 30 sessions from 4 mice | **X Seg 1:** 0.019<br>**X Seg 2:** 0.503<br>**Y Seg 1:** 0.015<br>**Y Seg 2:** 0.360<br>**Z Seg 1:** 0.903<br>**Z Seg 2:** 0.198 |
| 5M | Wilcoxon Signed-Rank Test | 30 sessions from 4 mice | **Seg 1:** 0.040<br>**Seg 2:** 0.221 |
| 5N | Wilcoxon Signed-Rank Test | 30 sessions from 4 mice | **Seg 1:** 0.004<br>**Seg 2:** 0.919 |
| 5O | Wilcoxon Signed-Rank Test | 30 sessions from 4 mice | **Left to Right (Lift to Retract):**<br>0.784 \| 0.382 \| 0.077 \| 0.005 \| 0.114 |
| 5P | Wilcoxon Signed-Rank Test | 30 sessions from 4 mice | **Left to Right (Lift to Retract):**<br>0.571 \| 0.584 \| 0.003 \| 0.008 \| 0.008 |
| S1B | Friedman test | 251 sessions from 28 mice | < 0.0001 |
| S1C | Friedman test | 251 sessions from 28 mice | < 0.0001 |
| S1D | Friedman test | 251 sessions from 28 mice | < 0.0001 |
| S1E | Friedman test | 251 sessions from 28 mice | < 0.0001 |
| S1F | Friedman test | 251 sessions from 28 mice | < 0.0001 |
| S1G | Friedman test | 251 sessions from 28 mice | < 0.0001 |
| S1H | Friedman test | 251 sessions from 28 mice | < 0.0001 |
| S1I | Friedman test | 251 sessions from 28 mice | < 0.0001 |
| S1J | Friedman test | 251 sessions from 28 mice | < 0.0001 |
| S1K | Friedman test | 251 sessions from 28 mice | < 0.0001 |
| S1L | Friedman test | 251 sessions from 28 mice | < 0.0001 |
| S1M | Friedman test | 251 sessions from 28 mice | < 0.0001 |
| S2B | Friedman test | 21 sessions from 7 mice | < 0.0001 |
| S2C | Friedman test | 21 sessions from 7 mice | < 0.0001 |
| S2D | Friedman test | 21 sessions from 7 mice | < 0.0001 |
| S2H | Friedman test | 19 sessions from 7 mice | < 0.0001 \| < 0.0001 \| < 0.0001 \| 0.0005 \| 0.065 |
| S2I | Friedman test | 19 sessions from 7 mice | 0.018 \| 0.0003 \| 0.0004 \| < 0.0001 \| 0.029 |
| S3C | Wilcoxon Signed-Rank Test | 14 sessions from 5 mice | **Within Laser:**<br>0.0001<br>**After Laser:**<br>0.0001 |
| S3F | Wilcoxon Signed-Rank Test | 14 sessions from 5 mice | 0.583 |
| S3G | Wilcoxon Signed-Rank Test | 14 sessions from 5 mice | 0.0004 |
| S3J | Wilcoxon Signed-Rank Test | 14 sessions from 5 mice | **Left to right:**<br>0.009 \| 0.007 \| 0.042 \| 1.000 \| 0.626 |
| S3K | Wilcoxon Signed-Rank Test | 14 sessions from 5 mice | **Left to right:**<br>0.013 \| 0.011 \| 0.013 \| 0.855 \| 0.626 |
| S4B | Wilcoxon Signed-Rank Test | 12 sessions from 4 mice | **20Hz - Control vs 40Hz - Control:** 0.301<br>**20 Hz vs Control:** 0.424<br>**40Hz vs Control:** 0.339 |
| S4C | Wilcoxon Signed-Rank Test | 12 sessions from 4 mice | **20Hz- Control vs 40Hz - Control:**<br>0.722<br>**20 Hz vs Control:**0.970<br>**40Hz vs Control:** 0.569 |
| S4D | Wilcoxon Signed-Rank Test | 12 sessions from 4 mice | **20Hz - Control vs 40Hz - Control** 0.233<br>**20 Hz vs Control:**0.910<br>**40Hz vs Control:** 0.110 |
| S4E | Wilcoxon Signed-Rank Test | 12 sessions from 4 mice | **20Hz - Control vs 40Hz - Control:** 0.077<br>**20 Hz vs Control:** 0.027<br>**40Hz vs Control:** 0.151 |
| S4F | Wilcoxon Signed-Rank Test | 12 sessions from 4 mice | **20Hz - Control vs 40Hz - Control:** 0.970<br>**20 Hz vs Control:** 0.001<br>**40Hz vs Control:** 0.007 |
| S4J | Wilcoxon Signed-Rank Test | 12 sessions from 4 mice | **20Hz- Control vs 40Hz-Control:**<br>0.233 \| 0.002 \| 0.677 \| 0.077 \| 0.204<br>**20 Hz vs Control:**<br>0.129 \| 0.034 \| 0.176 \| 0.077 \| 0.034<br>**40 Hz vs Control:**<br>0.064 \| 0.002 \| 0.110 \| 0.009 \| 0.009 |
| S4K | Wilcoxon Signed-Rank Test | 12 sessions from 4 mice | **20Hz- Control vs 40Hz-Control:**<br>1.000 \| 0.012 \| 0.110 \| 0.034 \| 0.034<br>**20 Hz vs Control:**<br>0.129 \| 0.012 \| 0.266 \| 0.064 \| 0.027<br>**40 Hz vs Control:**<br>0.424 \| 0.007 \| 0.002 \| 0.005 \| 0.002 |
| S5B | Wilcoxon Signed-Rank Test | 21 sessions from 7 mice | 0.881 |
| S5C | Wilcoxon Signed-Rank Test | 21 sessions from 7 mice | 0.838 |
| S5D | Wilcoxon Signed-Rank Test | 21 sessions from 7 mice | 0.683 |
| S5E | Wilcoxon Signed-Rank Test | 21 sessions from 7 mice | 0.157 |
| S6B | Wilcoxon Signed-Rank Test | 21 sessions from 7 mice | 0.708 |
| S6C | Wilcoxon Signed-Rank Test | 21 sessions from 7 mice | 0.0002 |
| S6D | Wilcoxon Signed-Rank Test | 21 sessions from 7 mice | 0.609 |
| S6E | Wilcoxon Signed-Rank Test | 21 sessions from 7 mice | 0.838 |
| S7D | Wilcoxon Signed-Rank Test | 26 sessions from 7 mice | **X Seg Mean:**<br>0.007 \| 0.143 \| 0.899 \| 0.216 \| 0.191<br>**Y Seg Mean:**<br>0.080 \| < 0.0001 \| 0.248 \| 0.027 \| 0.312<br>**Z Seg Mean:**<br>< 0.0001 \| < 0.0001 \| 0.002 \| 0.002 \| 0.120 |
| S7E | Wilcoxon Signed-Rank Test | 26 sessions from 7 mice | **Left to right:**<br>< 0.0001 \| < 0.0001 \| 0.085 \| 0.546 \| 0.634 |
| S8D | Wilcoxon Signed-Rank Test | 30 sessions from 4 mice | **X Seg Mean**<br>0.635 \| 0.038 \| 0.003 \| 0.324 \| 0.730<br>**Y Seg Mean**<br>0.898 \| 0.0005 \| 0.051 \| 0.324 \| 0.700<br>**Z Seg Mean**<br>0.689 \| 0.029 \| 0.701 \| 0.013 \| 0.038 |
| S8E | Wilcoxon Signed-Rank Test | 30 sessions from 4 mice | **Left to right:**<br>0.303 \| 0.019 \| 0.001 \| < 0.0001 \| 0.016 |

## Notes

### Competing Interest Statement

The authors have declared no competing interest.

### Summary of Updates

Minor formatting and spelling errors in figures have been corrected.

